# Global indicators of climate-driven shifts in vertebrate assemblages

**DOI:** 10.64898/2026.11.12.766169

**Authors:** Wendy B. Foden, Matthew J. Nagy-Watson, Vineet Katariya, Jeremy T. Kerr, Alan T.K. Lee, Steven R. Beissinger

**Author notes:** These authors contributed equally to this work (Wendy B. Foden, Matthew J. Nagy-Watson).

## Abstract

We develop global indicators for tracking how species assemblages respond to climate change by estimating community climate indices based on species’ maximum temperature and minimum aridity niche limits. We applied them to global databases using 33.6 million occurrence records from the past half-century to track assemblage changes for 26,163 species of terrestrial vertebrates. Changes in community climate indices tracked climate changes, weakening at higher elevations and strengthening at higher latitudes. Species that disappeared from sampled locations were intolerant of hot and dry conditions, while colonising species were characteristic of warmer and drier locales. Trends differed markedly within subcontinental regions, ecoregions and countries. Many vertebrate assemblages are responding to climate change yet lags in some regions and countries raise concerns that they are not keeping pace.

---

Climate change is causing global shifts in species’ distributions, phenology, morphology, and behaviour that may contribute to population declines and extinction (*1–5*). These impacts are widely recognised in global policy frameworks, including the Kunming-Montreal Global Biodiversity Framework under the UN Convention on Biological Diversity (CBD). Such policies anticipate the growing importance of climate change in the future, committing Parties to minimising climate-change impacts on biodiversity and building ecological resilience (Target 8) (*6*).

Delivering on these commitments requires robust indicators that enable countries to track how biodiversity is changing relative to climate change at spatial and temporal scales relevant to policy (*7*). Global indicators, such as the Red List Index (*8*), Living Planet Index (*9*, *10*) and Human Footprint Index (*11*), demonstrate the value of repeatable, publicly accessible metrics for tracking biodiversity. For climate change, the CBD endorses the Biodiversity Ecosystem Resilience Index (BERI), which measures a landscape’s capacity to promote or hinder climate-induced shifts in biological distributions (*12*), but it provides no direct measure of species’ responses. Studies that explicitly track climate change impacts on species (e.g., of European birds and butterflies (*13–16*); and global coral reefs (*17*, *18*)) have restricted regional and taxonomic focuses. The absence of a global, multi-taxa indicator for tracking biodiversity responses to climate change hampers measurement and reporting on progress for countries (*19*), complicating efforts to track and mitigate impacts on biodiversity.

Species are expected to respond directly to climate change because their physiological tolerances, performance optima, and demographic rates are constrained by environmental conditions, particularly temperature and water availability (*1*, *3*, *20*). As climates warm or become more arid, conditions at many locations approach or exceed niche limits of some species while becoming newly suitable for others, creating directional pressures on occupancy and persistence (*2*, *4*, *5*). These dynamics imply that climate change restructures communities through asymmetric processes of local colonisation and extirpation (*21*, *22*). Consequently, climate-driven biodiversity change should be detectable as systematic shifts in the climatic affinities of assemblages at a location, even where species richness remains stable (*15*, *23*). However, realised responses may change with differences in exposure due to microclimatic buffering and fine-scale habitat associations, causing assemblage composition to lag macroclimate trends (*23–26*). Indicators that summarise the climatic characteristics of assemblages provide mechanistically grounded and policy-relevant means of detecting and comparing real-world responses to climate change across space and time (*1*, *15*, *19*) rather than predicted responses from unverifiable models.

We develop a global indicator for tracking biodiversity responses to climate change using Community Climate Indices (CCIs) – specifically the Community Temperature Index (CTI) and the Community Aridity Index (CAI) – with open-access databases of species occurrences to show how vertebrate assemblages have changed over the past half-century. The CTI and CAI amalgamate species’ temperature and aridity affinities, respectively, and consolidate them into place-based measures for assemblages that directly reflect climate change and can be tracked over time (Fig. 1). Long-term monitoring has demonstrated that increases in the CTI of select assemblages correspond with increases in regional temperature (*13–15*, *27–29*).

**Fig. 1.**
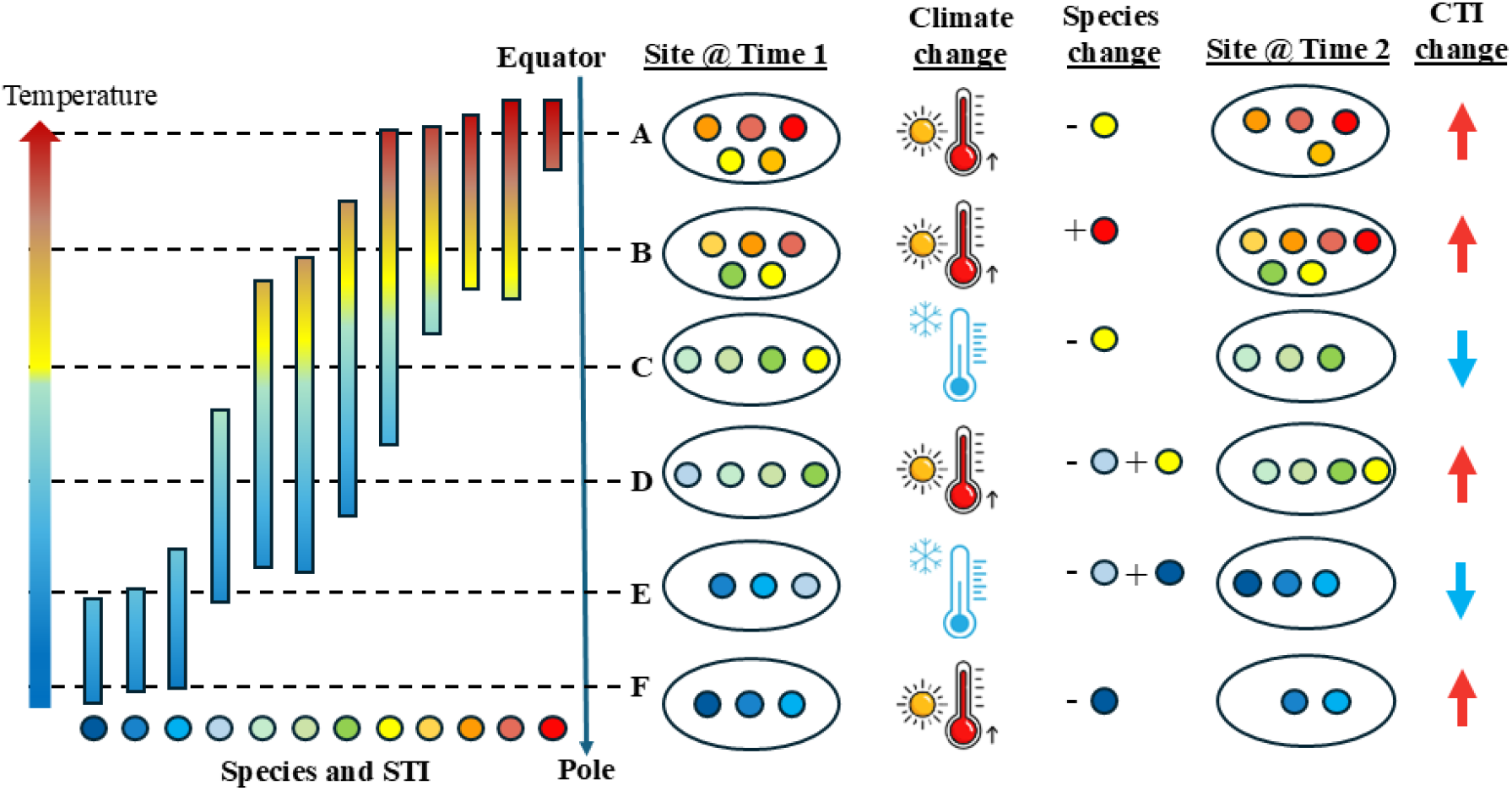
How the community thermal index (CTI) acts as an indicator of the effects of climate change on assemblages of species. Climate change causes species to track their thermal niches as sites change in suitability, which results in changes to species composition of assemblages and CTI. Species vary in the thermal niches they occupy, defined by the mean, upper and lower temperatures in their geographic ranges and quantified by the Species Temperature Index (STI). Species with ranges near the equator generally experience warmer environmental conditions and temperatures and have higher STIs (red = hot, blue = cold) than species found towards the poles. The CTI of an assemblage at a location is an emergent property of the climatic associations of its species and is calculated as the average of the STIs of each species at a site. We use maximum temperatures for calculating STI and CTI because climatic extremes set species range limits. Two processes related to climate change can lead to a change in the CTI of an assemblage over time: species arriving as a site becomes thermally suitable and species disappearing as it becomes unsuitable. Climate change has predominantly resulted in warming at most locations on Earth (sites A,B,D,F), but some sites have cooled (C,E). Climate warming results in the local extinction or disappearance (subtraction) of species with cooler STIs (sites A,D,F) and/or leads to the local colonisation or appearance (addition) of species with warmer STIs (sites B, D). Both changes result in an increase in the CTI. When a site cools, species with hotter STIs are likely to disappear (sites C, E) and may be replaced by species with cooler STIs (site E), resulting in a decrease of the assemblage’s CTI. When the hottest or coldest locations warm (site A) or cool (site F), respectively, beyond thermal niches, causing species to disappear, there may be no other species near the equator with hotter STIs or near the poles with colder STIs to replace them.

We applied these indices to global time series of terrestrial vertebrate occurrences to quantify shifts in species assemblage composition from 1960 to 2020 using three novel adaptations. First, to quantify assemblages and how their climatic affinities change over time, we use two global open-access datasets – geographic range data from the International Union for Conservation of Nature (IUCN) (*30*) to calculate the Species Temperature Index (STI) and locality records of occurrences from the Global Biodiversity Information Facility (GBIF) (*31*) to calculate CTI. Second, we similarly quantified CAI based on the Species Aridity Index (SAI) that reflects the degree of dryness at a location based on the ratio of precipitation to evapotranspiration (*26*). Third, climatic extremes test species’ tolerances and consequently exert greater effects on range limits than mean conditions (*32*, *33*). Therefore, we estimated species’ niche limits by calculating STI and CTI using annual maximum temperatures and by quantifying SAI and CAI using the annual minimum aridity index (see Supplementary Materials and Methods).

Here, we test the effectiveness of CCIs as indicators of biodiversity responses to climate change and evaluate their capacity to reflect climate change impacts on species assemblage change. Focusing on 26,163 vertebrate species (6,196 amphibians, 9,534 birds, 4,645 mammals, and 5,788 reptiles) and using >33.6 million occurrence records (Table S10), we hypothesised that species should respond to shifting climatic conditions and the resulting assemblages, and their CTI and CAI, would track local changes (Fig. 1). Specifically, in a warming and more arid world, we predicted that species associated with cooler (lower STI) and more mesic climates (higher SAI) would be preferentially lost from sites that warmed and dried, being replaced by species from warmer (higher STI) and more arid (lower SAI) climates. Finally, we expected response differences between vertebrate classes because they differ in thermoregulatory strategies, sensitivities to desiccation, and dispersal capacities, which alter their exposure, sensitivity, and adaptive capacity (*2*, *34*, *35*).

## Results

### Vertebrate assemblages are tracking climate change

CCIs tracked climate change in quadrats sampled across all decades (Fig. 2). The ambient global maximum temperature at sampled quadrats increased by approximately 1.37 °C from 1960-2020 (Fig. 2A). As expected, CTI for vertebrates also showed a clear increasing trend of approximately 0.348 °C over this period (Fig. 2C). Mean aridity index declined by approximately 0.078 from 1960-2020 (Fig. 2B), reflecting general drying trends over areas where terrestrial vertebrates were observed, and the CAI for all vertebrates exhibited a corresponding decline of approximately 0.012 over the same period (Fig. 2D).

**Fig. 2.**
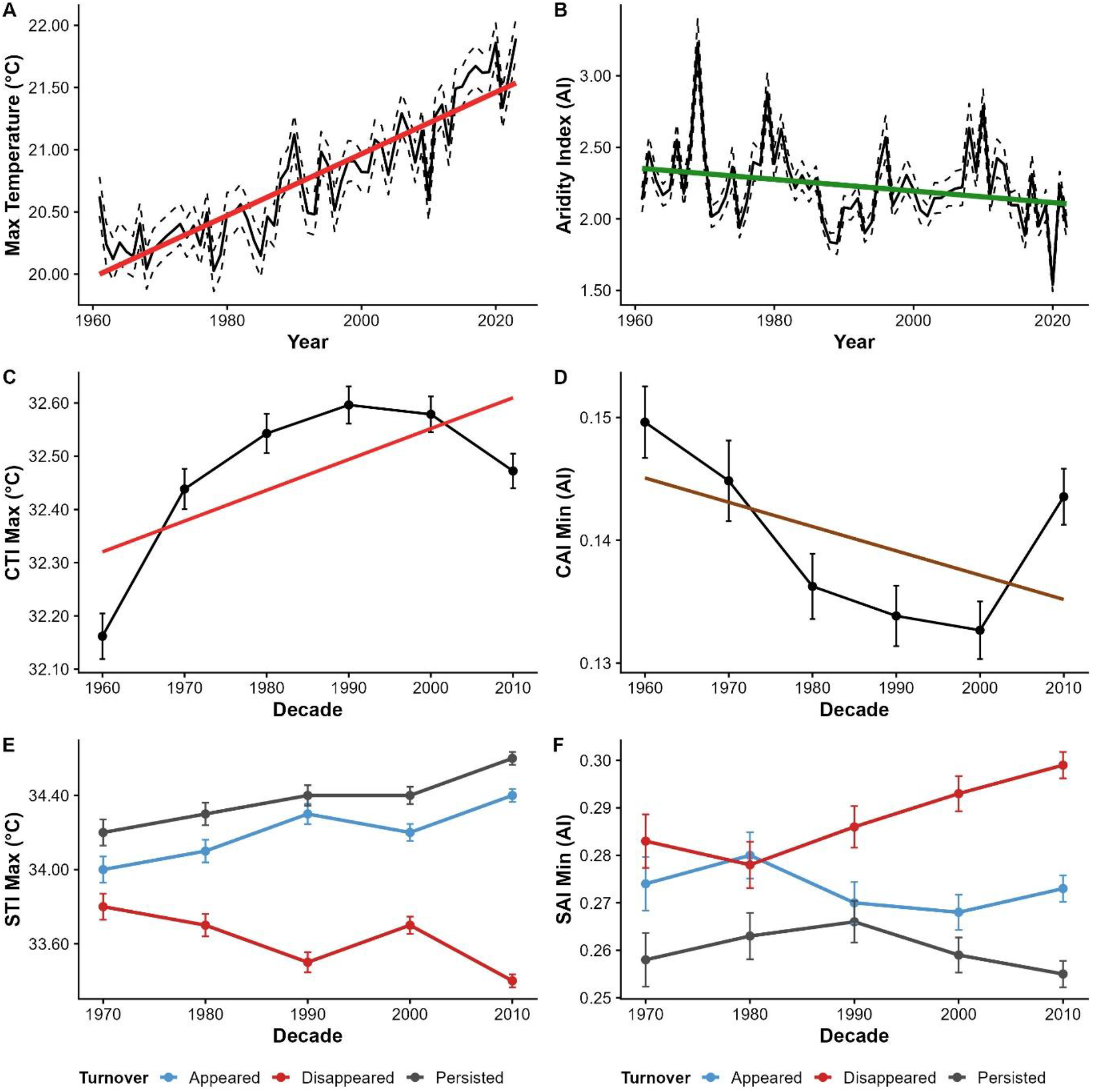
Temporal trends in climate and community temperature and aridity index in the study region. (**A**) Annual mean maximum temperature from 1960–2020, with the fitted linear trend shown in red, and (**B**) the Aridity Index over the same period, noting that decreasing AI values reflect drying. (**C**) Decadal changes in the Community Temperature Index (CTI) showing shifts toward warmer-affiliated communities through time (error bars indicate uncertainty around the mean, and the red line shows the long-term linear trend), and (**D**) decadal changes in the Community Aridity Index (CAI), indicating community drying over this period (error bars show uncertainty, and the brown line indicates the overall linear trend).

CTIs of each class of vertebrates increased from 1960-2020 for assemblages in the subset of quadrats sampled across all decades (Fig. 3A). Significant positive relationships between trends in average maximum temperature and CTI (Fig. 3B, Tables S1, S5) were strong for mammals and moderate for amphibians and reptiles; the relationship for birds was not significant. CTI trends were modulated by geography, weakening at higher elevations and strengthening at higher latitudes. CAIs for assemblages shifted less predictably, with mammals and reptiles declining, birds changing little, and amphibians increasing until declining in the final decade (Fig. 3D). Temporal trends in CAI (Fig. 3E, Tables S2, S5) were significantly positive for mammals and amphibians, but non-significant for birds and reptiles. Community aridity responses weakened significantly at higher elevations and latitudes.

**Fig 3.**
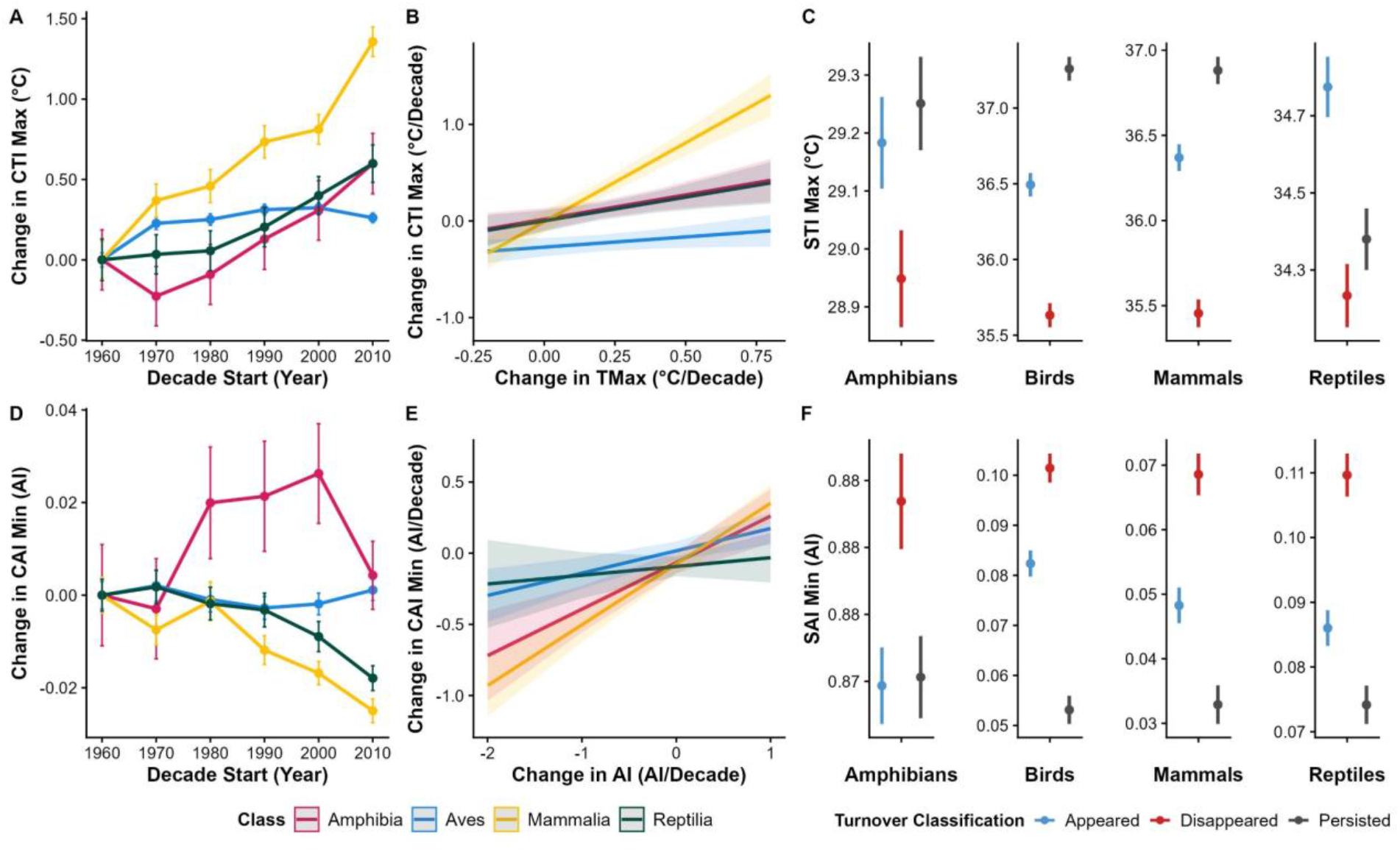
Temporal changes in community temperature (CTI) and moisture (CAI) indices across vertebrate groups. (**A**) Decadal change in Community Temperature Index (CTI) from 1960–2010, showing increasing thermal affinity through time, with coloured lines representing major vertebrate groups (amphibians, birds, mammals, reptiles) and dashed lines indicating uncertainty. (**B**) Relationship between CTI trends and maximum temperature (TMax) trends, with fitted lines and confidence intervals by taxonomic group. (**C**) Mean STI of species that appeared, disappeared, or persisted within assemblages, separated by taxonomic group; points show estimates with uncertainty. (**D**) Decadal change in Community Aridity Index (CAI) over the same period, showing group-specific responses to changing moisture conditions. (**E**) The relationship between CAI trends and aridity index (AI) trends, showing how assemblage moisture affinity tracks climatic drying or wetting. (**F**) mean SAI of species that appeared, disappeared, or persisted, highlighting that newly appearing species tend to be more warm- and dry-adapted than those that disappear.

### Vertebrate assemblage changes were driven by climate filtering

To investigate the processes underlying CCI trends, we compared STIs and SAIs of species that disappeared, appeared, and persisted in quadrats. We anticipated the disappearance of cool-preferring species (lower STIs) and the appearance of warm-preferring species (higher STIs) because of long-term warming (Fig. 2A). Our findings confirmed this trend for global terrestrial vertebrates (Fig. 2E) and for all four vertebrate classes (Fig. 3C, Tables S3, S6). The difference in STI between appearing and disappearing species widened from 1960-2020 for all vertebrates (Fig. 2E). STIs for species that persisted were also higher than species that disappeared for all vertebrates (Fig. 2E) and for all classes (Fig. 3C, Tables S3, S6). Species that persisted had the highest STIs for all classes except reptiles. These results reveal a consistent signal of climate-driven thermal filtering.

We anticipated that wet-affiliated species (higher SAIs) would preferentially disappear and arid-tolerant species (lower SAIs) would appear, given the overall drying trend we detected (Fig. 2B). Our findings confirmed this prediction: disappearing species had higher SAIs than appearing species for all vertebrates in all decades except for 1970-1980 (Fig. 2F), and the gap between them widened successively from 1980 to 2020 (Fig. 2F). Species that persisted had lower SAI than species that disappeared or appeared (Fig. 2F). These patterns held for each vertebrate class except amphibians, where the SAIs of persisting and appearing species were similar (Fig. 3F, Tables S4, S7). Unsurprisingly, amphibians are strictly tied to high-moisture/low-aridity, whereas other classes were less constrained. These results confirm a climate-driven aridity filtering effect across terrestrial vertebrates.

### Hotspots of vertebrate assemblage change

Globally, all regions experienced both increases and decreases in CTI and CAI (Fig. 4C,D) in response to climatic changes (Fig. 4A,B), but clear trends emerged. Vertebrate CCIs for Western Europe, the southern Andes and the Himalayas underwent assemblage warming and wetting, while Eastern Europe, northern Scandinavia, south-eastern Australia, and New Zealand experienced assemblage warming and drying. Assemblages that predominantly cooled and became wetter include the Indian subcontinent, southern Scandinavia, South-East Asia, south-eastern Africa, and central Australia. A few areas showed mainly assemblage cooling and drying, including parts of central North America, western Asia and southwestern Australia.

**Fig 4.**
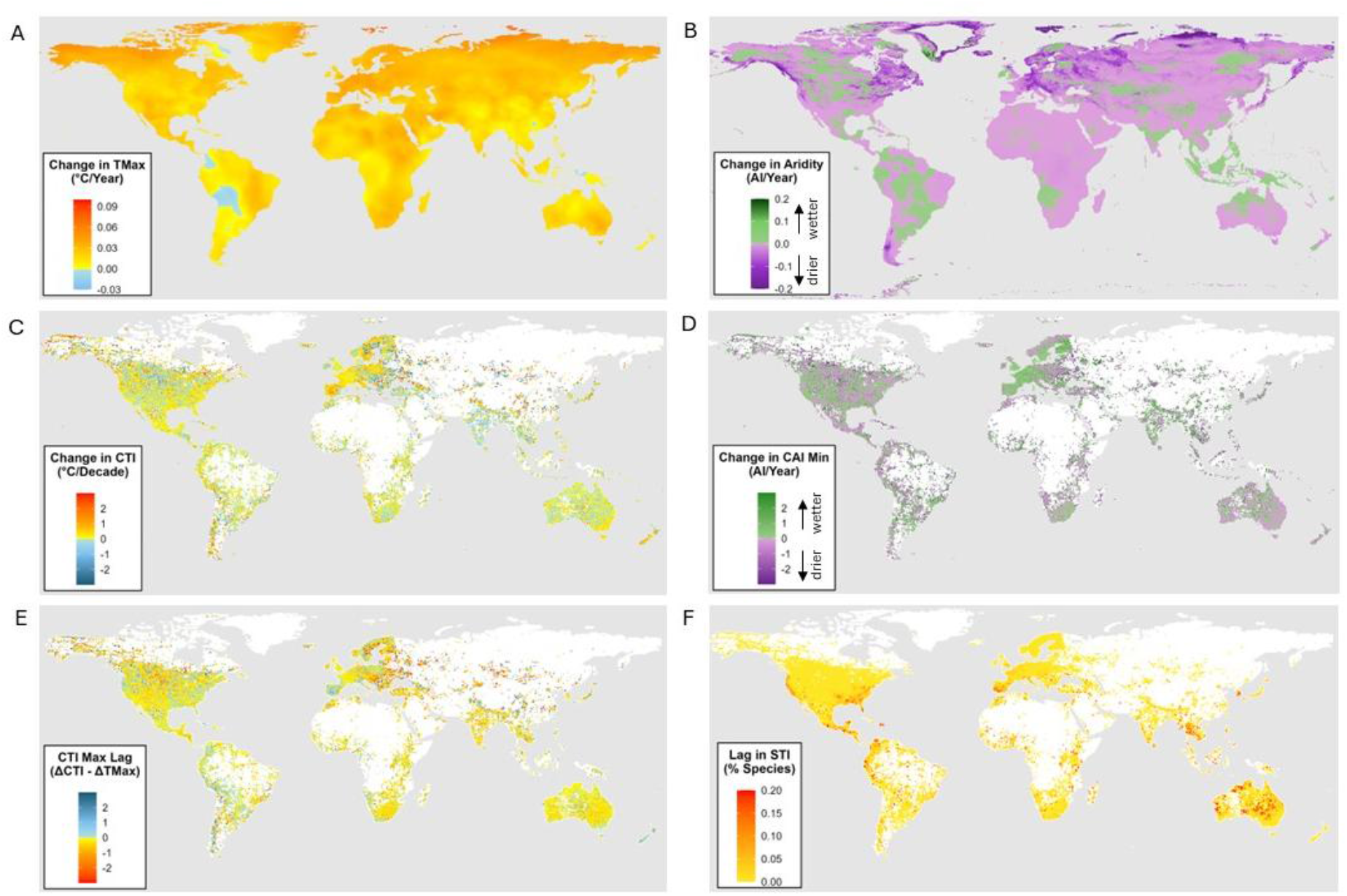
Global patterns of warming and wetting, and the extent to which communities have tracked them. Maps show regional patterns of change in (**A**) maximum temperature (°C/yr) and (**B**) the Aridity Index (AI/year) from 1960-2020. Decadal trends in (**C**) the Community Temperature Index (°C/decade) highlight where species assemblages are becoming more warm- or cool-affiliated, and those of the (**D**) Community Aridity Index (AI/decade) indicate shifts toward more dry- or wet-adapted assemblages. The lag in assemblage reassembly (**E**) is represented as the difference between maximum temperature (Tmax) change and CTI change (1960-2020). The lag in STI (**F**) shows the percentage of species for which the average Tmax over the last decade exceeds their STI, reflecting species for which maximum temperature thresholds may already have been exceeded.

CTIs of vertebrate assemblages in most ecoregions warmed significantly (Figs. 5A, S2, Table S8), like their underlying climates (Fig. S4), but poleward ecoregions changed faster than equatorial regions, led by Tundra and Boreal Forests (Taiga). Tropical and subtropical forest grassland and shrubland ecoregions show the least CTI change, along with Desert ecoregions. There are few species with warmer climate niches to replace the ones that currently inhabit these regions as they warm. CAI change was less common (Figs. 5B, S3, Table S9), with 10 of 16 ecoregions exhibiting nonsignificant assemblage shifts as aridity trends varied greatly among ecoregions (Fig. S5). Vertebrate assemblages of Tundra and Boreal Forests/Taiga ecoregions dried the most, followed by Temperate Conifer and Montane Grasslands. No ecoregion assemblages became occupied by wetter species.

**Fig. 5.**
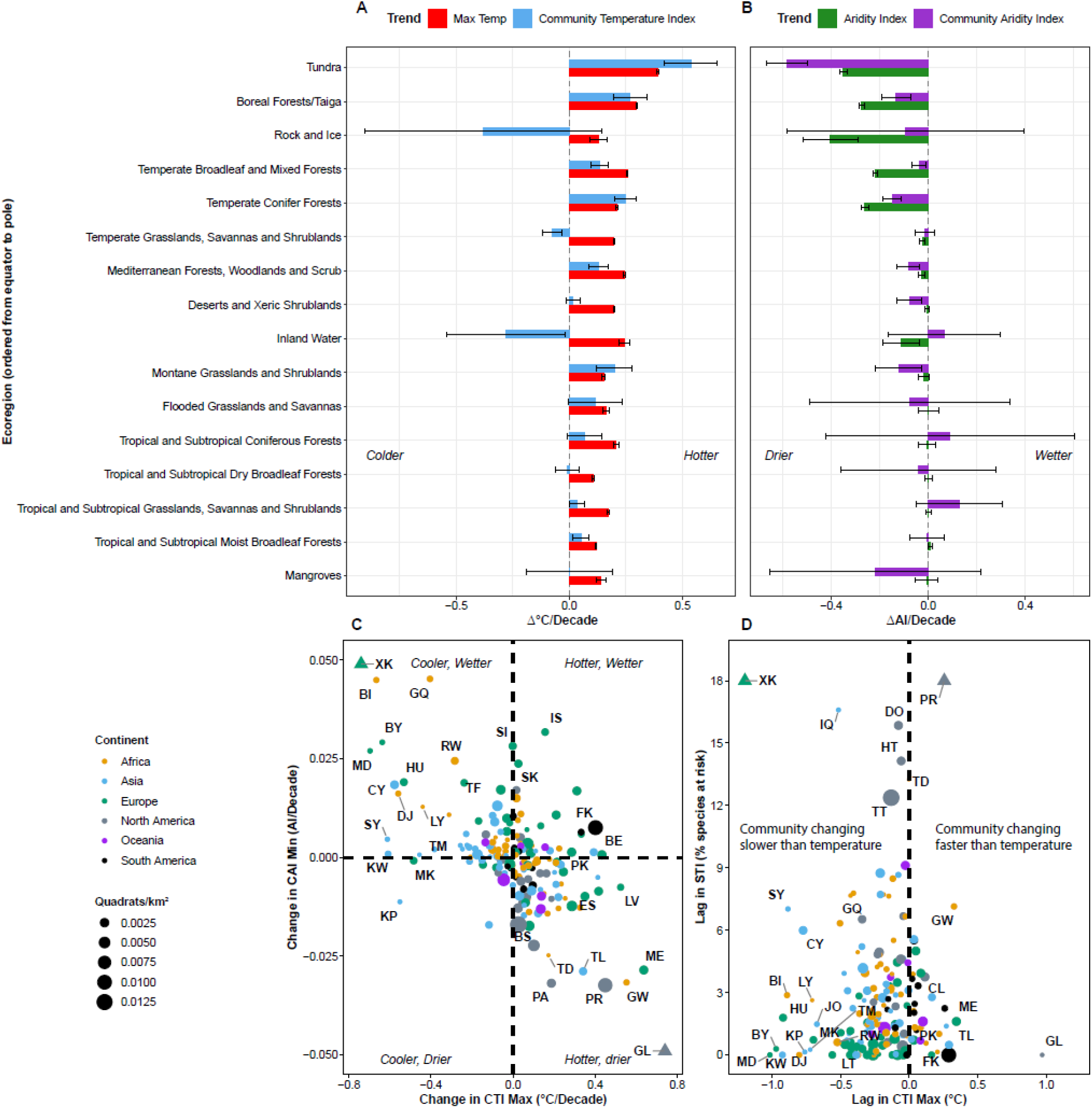
Country-level changes of community climate indices for terrestrial vertebrate assemblages and their thermal risk. (**A**) Change in the CTI maximum and CAI minimum by country (1960-2020). Triangles indicate two outliers: Greenland (GL) (x = 1.13, y = -0.10), and Kosovo (XK) (x = -2.60, y = 0.04). (**B**) Thermal risk to vertebrate assemblages is represented on the X-axis as the lag in change between CTI relative to change in maximum temperature, averaged per country. The Y axis shows the lag in STI, the percentage of species for which the average Tmax over the last decade exceeds their STI, reflecting species for which maximum temperature thresholds may already have been exceeded. Dot size is scaled by sampling coverage and coloured by continent. Triangles indicate outliers Kosovo (XK) (-2.88, 0.20) and Puerto Rico (PR) (0.26, 0.24). Two letter country codes are: BE = Belgium, BI = Burundi, BS = Bahamas, BY = Belarus, CL = Chile, CY = Cyprus, DJ = Djibouti, DO = Dominican Republic, ES = Spain, FK = Falkland Islands, GL = Greenland, GQ = Equatorial Guinea, GT = Guatemala, GW = Guinea Bissau, HT = Haiti, HU = Hungary, IQ = Iraq, IS = Iceland, JO = Jordan, KP = North Korea, KW = Kuwait, LT = Lithuania, LV = Latvia, LY = Libia, MD = Moldavia, ME = Montenegro, MK = North Macedonia, PA = Panama, PK = Pakistan, PR = Puerto Rico, RW = Rwanda, SI = Slovenia, SK = Slovakia, SY = Syria, TD = Chad, TL = Timar-Leste, TM = Turkmenistan, and XK = Kosovo.

Climate change pulled countries’ vertebrate assemblages in opposing directions over the past 50 years (Fig. 5B). Assemblages of 37.5% of countries became hotter and drier, while 30.7% became cooler and wetter. Only 11.3% of assemblages became cooler and drier, while the remainder (20.5%) got hotter and wetter. Countries dominated by assemblages that increased in temperature and aridity included Greenland, Guinea-Bissau, Puerto Rico, Montenegro, Panama, and Timor-Leste, while assemblages that cooled and became more mesic included Kosovo, Burundi, Equatorial Guinea, Belarus, Cyprus, and Moldova.

### Vertebrate assemblage changes often lagged climate change

The relationship between changes in CTI and changes in maximum temperature quantifies whether assemblages are keeping pace with thermal change and reveals substantial geographic variation (Fig. 4E). The largest lags occurred in Eastern Europe, Scandinavia, parts of central Asia and the Andes, the southern United States, southern Africa, and much of the Indian subcontinent, Southeast Asia, and Australia. In contrast, some regions exhibited rates of reassembly that matched or exceeded ambient warming, including western Europe, western and eastern Australia, and parts of central and western South America. Western Europe provides a notable example of rapid community adjustment. Despite experiencing substantial warming and large increases in CTI, the region exhibited relatively small lags because assemblage change closely tracked temperature changes. An equivalent measure of lag for CAI minimum is not possible to estimate because temporal trends in CAI based on the minimum value in a species’ range were ∼10 times smaller than mean AI trends. This generated a lag metric that was near-perfectly correlated with AI trends.

At the country scale, assemblage change lagged climate change in 82.3% of countries, with 10% exhibiting strong lags (< 0.5 °C) (Fig. 5B). The largest lags occurred in Kosovo, Belarus, Hungary, Kuwait, Syria, Cyprus, North Macedonia, and Libya. Greenland was the only country where assemblage change strongly outpaced climate warming, as primarily local species expanded their ranges and abundances to locations that recently became ice-free (*36*).

### Where warming exceeds species tolerances

We evaluated thermal risk by comparing the lag in CTI to the percentage of each quadrat’s species for which critical temperature thresholds may have already been exceeded over the last decade. These species represent a thermal extinction debt because they are likely to experience reduced reproduction and survival, and are expected to disappear in the absence of mechanisms that reduce their thermal vulnerability. Globally, areas of the highest species risk (Fig. 4F) include Western Europe, large parts of central and western Australia, Southeast Asia, coastal and eastern North America, Central America, northwestern South America and parts of Africa. Countries with assemblages at most thermal risk (12-24%) (Fig. 5D) include Kosovo, Puerto Rico, Iraq, Dominican Republic, Haiti, and Chad. A second tier of risk includes countries with 5-10% lags in STI and lags in CTI > 0.5°C (Burkina Faso, Cambodia, Eq. Guinea, Eritrea, Guinea, Honduras, Madagascar, Myanmar, N. Cyprus, Niger, Somalia, South Korea, Syria, and Thailand).

## Discussion

### Vertebrate assemblage responses to climate change

Our results demonstrate direct, climate-associated community reassembly over the past half-century based on observed biodiversity responses. CCI changes reveal turnover in vertebrate assemblages, driven by climate change, that follows expectations derived from the indicators we propose (Fig. 1). STIs and SAIs generally show species that disappeared from quadrats are intolerant of hot and dry conditions, and species that appeared are characteristic of warmer and drier locations (Figs. 2E,F, 3C,F). Moreover, species that persisted in quadrats throughout the study’s six decades were generally the most heat- and arid-tolerant of all.

Vertebrate classes exhibited consistent responses to climate change, but some differences emerged (Fig. 3). CTIs of all classes but birds increased significantly. Birds had the highest STIs (Fig. 3C), suggesting exposure to climate warming may have exceeded avian thermal niches less frequently than for other taxa. Moreover, birds are the most mobile, enabling them to buffer against warming by exploiting fine-scale thermal heterogeneity resulting from aspect and canopy cover (*25*), although these strategies are not always effective (*24*, *37*, *38*). CAI trends exhibited greater variation among vertebrate classes, with the largest changes in mammals and amphibians and weaker responses in reptiles and birds (Fig. 3D,E). Amphibians are directly constrained by moisture availability (*39*, *40*), while mammals often depend on vegetation productivity and surface water closely linked to aridity metrics (*41*, *42*), explaining the strong aridity relationships observed for these groups. Many reptiles are well adapted to arid environments, possessing physiological traits and behavioural strategies to minimise evaporative water loss (*3*, *43*). Although aridity can drive avian turnover directly through physiological water stress in hot environments (*24*, *37*, *38*), it may frequently affect avian assemblage turnover indirectly via changes in food availability, vegetation, or breeding success (*44*). Consequently, increasing aridity may not lead to rapid assemblage change in birds over decadal timescales.

While CCI changes that track climate changes indicate assemblage adaptation, the implications of lagged or seemingly decoupled responses, which occurred at subcontinental, ecoregional and country scales (Figs. 4,5), are less clear. One explanation is that immigration is not keeping pace with environmental change (Table S12), and communities are accumulating extinction debt (*21*, *22*), with likely negative conservation outcomes. Alternatively, lagged responses may reflect a conservation-positive context of low climate sensitivity or high adaptive capacity: assemblage change is unnecessary because species can persist *in situ* through microhabitat buffering (*23*, *25*, *26*), phenotypic plasticity (*45*, *46*), behavioural plasticity (*47*), dispersal to higher elevations, or evolutionary change (*20*, *48*– *50*). In some regions, the failure of CCIs to track climate may signal that non-climatic pressures dominate community impacts, such as habitat loss, disease or other threats. Methodological and data limitations may also contribute to apparent mismatches between climate and community change. Limitations include the omission of climatic drivers beyond temperature and aridity, exclusion of species-poor quadrats, spatial biases in occurrence records, imperfect detection, and uncertainty in historical observations (Table S12).

Ambiguities in the mechanisms responsible for strong lags limit the diagnostic utility of these indices for conservation. Where climate is changing substantially and lags are low, however, conservation implications are clearer, tentatively suggesting that communities are adapting to climate change. Monitoring over time will indicate whether CCIs in lagging regions recouple or further decouple from their climate trends and, in combination with global databases of biodiversity change, should clarify the nature of lags. Meanwhile, conservation approaches that promote species movement (e.g., enhancing habitat availability and connectivity, assisting colonisation, and reducing non-climatic threats) will promote adaptive community reassembly in response to climate change.

### CCIs as global indicators of climate change impacts on biodiversity

We present a globally and locally scalable indicator for tracking responses of biological communities to climate change, enabling comparison of contemporary assemblages against historical baselines. CCI trends can be tracked for nations and ecosystems from regional to continental scales. Combining two open-access datasets, GBIF and IUCN Red List range maps, enables the application of the CCIs at a global scale. Although spatial coverage of GBIF data remains limited in some high-biodiversity areas of Africa, South America, Asia, and the Middle East, continued growth in GBIF records (including from citizen science), advances in AI-assisted species detection, eDNA, remote sensing and other monitoring technologies will further improve the spatial and temporal resolution of biodiversity observations, supporting regular CCI updates. Expanding data coverage, especially for plants and invertebrates, is essential.

CCIs have clear policy relevance. Nations can use the indicators to develop and report progress toward national biodiversity and climate commitments. Global bodies, including IPBES and the IPCC, can draw on them to communicate biodiversity change in a warming world. In addition to tracking climate change impacts (Target 8), the CCIs contribute to monitoring other Kunming-Montreal Global Biodiversity Framework targets, including preventing extinctions (Target 4). CCIs also offer opportunities for validating modelled range-shift projections, both past and future, including those served publicly (e.g., WildCRU (*51*) and SDMapCH (*52*)) and used in the Biodiversity Ecosystem Resilience Index (BERI) (*12*). Moreover, CCIs are simple to explain, straightforward to implement, compatible with existing global data infrastructures, and supported by robust, continually updated datasets of species occurrences. Previous endorsement of CTI-based indicators under the CBD further facilitates the potential for mainstream adoption.

## Conclusions

We have demonstrated the utility of CCIs based on freely available datasets for tracking species assemblage responses globally to climate change. Our results provide reassurance that many vertebrate assemblages are responding to climatic change, yet strong lags in some countries and regions raise concerns that assemblage change is not keeping pace. Coordinated policy action, increased conservation investment, enhanced monitoring and expanded data coverage are urgently needed. Critically, even modest reductions in the rate of climate change will help ecological communities re-align with shifting environmental conditions.

## Acknowledgments

Zishan Ebrahim assisted with maps, and Neil Burgess provided comments on the manuscript.

## Funding

Benjamin Raymond Oppenheimer (BRO) Trust (WBF)

United States Department of State grant SSF75022CA0040 (WBF)

Caldwell First Nation (MJN-W)

Discovery Grant and Discovery Accelerator support from the Natural Sciences and Engineering Research Council of Canada (JTK (grant holder) and MJN-W)

Mapula Trust (ATKL)

## Author contributions

Conceptualisation: WBF, MJN-W, ATKL, SRB

Methodology: WBF, MJN-W, VK, JTK, ATKL, SRB

Analyses: MJN-W, SRB Coordination: WBF

Figures: MJN-W, SRB, WBF

Writing – original draft: WBF, MJN-W, SRB, JTK, ATKL

Writing – review & editing: SRB, WBF, MJN-W, JTK, ATKL

## Diversity, equity, ethics, and inclusion

MJN-W identifies as Anishinaabe and is a member of Caldwell First Nation.

## Data, code, and materials availability

All data are available in the main text or the supplementary materials.

## Supplementary Materials

Materials and Methods

Supplementary Text

Figs. S1 to S5

Tables S1 to S13

Data S1 to S4

References *(53-69)*

