## Supplementary Information for "Global indicators of climate-driven shifts in vertebrate assemblages"

Corresponding author: Wendy Foden

**The PDF file includes:**

Materials and Methods
Supplementary Text
Figs. S1 to S5
Tables S1 to S13
Data S1 to S4
References

### Materials and Methods

<u>Step 1: Calculating Species Temperature and Aridity Indices</u>

*1.1 Preparing species distribution data*

Monthly thermal and AI niche limits were calculated to account for phenotypic plasticity that can allow for seasonal variability in species' tolerance to environmental conditions. Expert-informed range maps, as well as historical observations of species occurrences, can be used to generate species-level environmental estimates. While biases exist for both methods (Araujo & Peterson 2012), at global spatial scales, range maps and species observations generate similar species-level environmental estimates (Sunday et al., 2012; Khaliq et al., 2014). Since observations are often unevenly sampled across taxa (Beck et al. 2014), and do not cover as many species as is possible with expert-informed range maps, we decided to use range maps for deriving lower and upper thermal and aridity niche limits.

Following the download of global climate data, we accessed expert-informed range maps for all terrestrial amphibians, birds, mammals, and reptiles (IUCN, 2023; BirdLife International, 2023). Range maps included metadata describing birds' seasonal distributions, including resident regions (species are known to occur year-round), breeding season, non-breeding season, and passage (migratory regions). We generated four distinct seasonal range maps for all bird species: Breeding (resident and breeding range maps), Non-Breeding (resident and non-breeding range maps), Migration Non-Breeding (resident, non-breeding, and passage range maps), and Migration Breeding (resident, breeding, and passage range maps). To account for differences in the seasonal timing of migration and breeding periods between the Northern and Southern Hemisphere we classified all bird species distributions as Northern or Southern using biogeographic classifications (IUCN, 2022). Based on the hemisphere classification, we assigned range maps to corresponding months of yearly migration and reproduction cycles (Table S1). Range maps for amphibians, mammals, and reptiles had no seasonal classification, so we used the complete species range maps for every month in the downstream data extraction methods.

*1.2 Estimating species' climatic limits*

We chose the time-period 1961-1975 to represent historical environmental conditions that species would have experienced prior to rapid changes in global climate. We recognize that species' potential to adapt to changing environmental conditions, but note that current rates of climate change are estimated to greatly exceed historical rates of niche evolution (Quintero and Wiens 2013) and that vertebrates' generally relatively slow reproductive cycles reduce the likelihood of rapid evolutionary adaptation. We therefore assume that species' thermal and aridity niche limits are likely not evolving in pace with climate change and may be calculated using data prior to rapid climate change. This reduces the risk of overestimating species' environmental niche limits. Additionally, setting a baseline period for generating species' environmental niche limits allowed us to evaluate the temporal effects of climate change on species' relative thermal and aridity niche positions.

*1.2.1 Preparing climate data*

We accessed historical (1961-2023) monthly maximum ($T_{max}$) and minimum ($T_{min}$) temperature raster datasets from the WorldClim database (Fick & Hijmans, 2017), and monthly precipitation and potential evapotranspiration raster datasets from the TerraClimate database (Abatzoglou et al., 2018) at a spatial resolution of 2.5 minutes (~21 $km^2$ at the equator). To generate Aridity Index (AI) datasets, we divided monthly precipitation by potential evapotranspiration for every month between 1961-2023. We additionally generated monthly mean temperature ($T_{mean}$) datasets by averaging each grid cell between $T_{max}$ and $T_{min}$. Next, we aggregated and averaged $T_{max}$, $T_{min}$, $T_{mean}$, and Aridity Index (AI) to the same spatial resolution as our quadrats (50km x 50km) using the same historical climate datasets (WorldClim and TerraClimate).

*1.2.2 Calculating Species Temperature and Aridity Indices (STI and SAI)*

Using species range maps, we subset monthly global $T_{max}$, $T_{min}$, and AI to each species range and extracted the maximum and minimum values for each month from 1961 to 1975. We additionally extracted monthly mean range-wide $T_{mean}$ and AI. Global $T_{mean}$ raster files were generated by taking the mean of monthly $T_{max}$ and $T_{min}$ for each quadrat for each month-by-year combination. For birds, we determined which range map was used for the extraction process based on the seasonal classification (Table S1). We then generated mean, lower and upper thermal and AI niche limits for each month. Finally, we averaged monthly means for each niche limit variable across all months, resulting in three species temperature indices ($STI_{max}$, $STI_{mean}$ and $STI_{min}$) and three species aridity indices ($SAI_{max}$, $SAI_{mean}$ and $SAI_{min}$).

### **Step 2: Calculating Community Temperature and Aridity Indices (CTI and CAI)**

*2.1 Preparing species distribution data*

Observation records for all species of terrestrial vertebrates (amphibians, birds, mammals, and reptiles) were downloaded from the Global Biodiversity Information Facility (GBIF) for the years 1961-1975 (Tables S11, S13). We also downloaded all terrestrial vertebrate observations from the years 1976-2023 for species that were present in the initial dataset (1961-1975). This reduced the potential impact of increasing sampling intensity over time, which would otherwise inflate the number of new species detections. After all records were downloaded, we harmonised species names using the *taxize* (0.10.0) package in R to correct misspelt or out-of-use species binomials (Chamberlain & Szöcs, 2013; Chamberlain *et al.*, 2020). We then filtered all observations that lacked or contained impossible geographic coordinates and assigned observations to the decade of observation (1961-1969, 1970-1979, 1980-1989, 1990-1999, 2000-2009, 2010-2023). We also generated 50km-by-50km gridded quadrats at a global scale. Grouping observations by decade and quadrat of occurrence, we then condensed observations to a single occurrence for each unique species binomial per quadrat. This gave us records of all species occurrences within each quadrat over time. Finally, species for which names could not be resolved to match those of the IUCN range maps used to generate STI/SAI metrics in Step 1 were removed.

*2.2 Calculating Community Temperature and Aridity Indices (CTI and CAI)*

After extracting climate data for all observations, we added the previously derived thermal and aridity niche limits corresponding to each unique species observation. For each quadrat-by-decade group, we generated community upper ($CTI_{max}$), lower ($CTI_{min}$), and mean ($CTI_{mean}$) thermal niche limits, along with upper ($CAI_{max}$), lower ($CAI_{min}$), and mean ($CAI_{mean}$) aridity limits, by averaging thermal and aridity niche estimators for all species that were present.

We set a threshold minimum of five unique species occurrences, removing all quadrat-by-decade combinations that contained fewer unique species observations. We used this threshold to ensure that community estimates in downstream analyses contained sufficient records to be informative and were not overly sensitive to the loss or inclusion of a single species. We tested additional thresholds of two and ten unique species occurrences, and the results for ten unique species matched those using a threshold of five. Models were mostly consistent using the two unique species thresholds, but there were slight differences in model estimates. Since our threshold of five unique species is more robust to the gain or loss of smaller numbers of species and aligns with results generated from analyses using a ten-species threshold, we report the results from those analyses. Finally, we removed all quadrats that had observations for only a single decade.

For each quadrat, we then linearly regressed each community metric by decade to estimate the temporal trends in these metrics. For each quadrat, we extracted the slope coefficient of each trend, as well as the standard error estimates. We then repeated all calculations of CTI and CAI, as well as temporal trends in these indices, for each taxonomic class separately, to test for potential differences in responses across taxa in downstream analyses.

Generally, our method for estimating $CTI_{mean}$ and $CAI_{mean}$ indices is comparable to the Community Temperature Index (CTI) (Devictor et al., 2008). However, where CTI uses mean annual range-wide temperature like our generation of $CTI_{mean}$, our method includes separate indices for species upper ($CTI_{max}$) and lower ($CTI_{min}$) niche limits. Including measures of species upper and lower niche limits has been shown to predict species occupancy and physiological sensitivity to climate change (Sunday et al, 2014; Huey et al., 2012). Specifically, these estimates provide an indication of differences in species' ability to cope with extremes of temperature and aridity, which are not detectable using CTI/$CTI_{mean}$. This is due to the potential for species to have similar range-wide mean environmental conditions but different range-wide extremes. Further, we expand community environmental indices to include metrics of AI. Aridity levels indicate overall water availability and primary productivity, which are vital for species survival. Since global AI conditions are being altered by climate change, being able to identify and predict which species and communities are at greatest risk is vital.

### Step 3: Analyze Trends and Patterns

#### *3.1 Exploring vertebrate trends globally*

To test if CTI and CAI trends responded to climate change, we used a linear mixed effects model (LMM) approach. We generated separate models for each community metric ($CTI_{max}$, $CTI_{min}$, $CTI_{mean}$, $CAI_{max}$, $CAI_{min}$, and $CAI_{mean}$), with the temporal trend of each metric serving as the response variable. Our predictor variables included the temporal trend in $T_{max}$ for the $CTI_{max}$ model, $T_{min}$ for the $CTI_{min}$ model, $T_{mean}$ for the $CTI_{mean}$ model, and AI for all CAL models. Additionally, for all models we included mean elevation and Latitude as a second-order polynomial based on the center of each quadrat. We additionally included the interaction between climate trend variables and both latitude and elevation. Latitude and elevation terms were included to control for broad geographic differences in climatic trends and conditions between quadrats. Due to evolutionary and physiological differences between classes of terrestrial vertebrates that could impact community responses to climate change, we included taxonomic class and the interaction between class and climate trend in all models. Landscape characteristics and ecosystem composition can impact exposure to prevailing climatic conditions. We accounted for differences in these ecosystem characteristics by including quadrat ecoregion as a random intercept variable. Finally, we included a weighting term in each model to account for uncertainty in measurements of CTI and CAI trends. The weighting term was calculated as the $\log_{10}$ value of the inverse standard error for each CTI and CAI slope coefficient generated during the temporal trend modeling step outlined above. All models were tested and found to conform to the assumptions of linear mixed effects models. Using spline correlograms, we found no evidence for spatial autocorrelation in model residuals.

Following the general methods in Lajeunesse and Fourcade (2022), for every month, we linearly regressed each environmental variable by year for every grid cell. We then averaged all trends for each quadrat to get climatic trends for $T_{max}$, $T_{min}$, $T_{mean}$, and AI for each quadrat. Additionally, we averaged all environmental variables for each decade to get mean $T_{max}$, $T_{min}$, $T_{mean}$, and AI values for each quadrat. For each species observation in the GBIF dataset, we extracted the temporal trends of $T_{max}$, $T_{min}$, $T_{mean}$, and AI based on quadrat, and mean decadal $T_{max}$, $T_{min}$, $T_{mean}$, and AI based on quadrat and decade. Additionally, we calculated mean elevation (European Space Agency, 2024) and extracted the most common ecoregion (Olson, 2020) for each quadrat and added these values to the dataset.

#### *3.2 Exploring trends taxonomically*

Next, we explored the processes underlying changes in CTI and CAI by testing if STI and SAI indices differed between species that disappeared from a community and those that appeared or persisted. We started by generating turnover classifications for each species within a given quadrat and decade. Within each quadrat, we classed species as "disappeared" if they were observed in a given decade but not in the immediately following decade. Next, we classified all species as "persisted" if they were observed in a given decade and the following decade. Finally, species were classed as "appeared" if they were present in a given decade but were not observed in the previous decade. We then grouped all species by quadrat, decade, and turnover classification and calculated all three STI and SAI indices. Next, we developed separate linear mixed effects models for each variable, six in total. Each model contained an STI or SAI index as the response variable. Then we included turnover classification as a predictor variable. For the same reason as our previous analyses, we included mean quadrat elevation and latitude as a second-order polynomial based on the centre of each quadrat. Due to this dataset containing repeated measures within quadrats and decades, we included both terms as random intercept variables. We also included taxonomic class and its interaction with turnover classification to identify if variable responses between classes of terrestrial vertebrates existed. To account for the influence of the number of species observations potentially influencing model results, we included the number of species as a weighting term. All models were evaluated to ensure the assumptions of linear mixed effect models were met. Using spline correlograms, we found no evidence for spatial autocorrelation in model residuals.

#### *3.3 Exploring trends spatially*

To test differences among ecoregions, we used the same model structure as used for investigating differences in CTI and CAI trends, but ecoregion was elevated to a fixed effect. However, CAI models with an ecoregion fixed effect and a random effect for quadrats yielded a singular fit, caused by parameters estimated on the boundary of the feasible space, so the random effect was removed, yielding a valid general linear model.

To generate plots of temporal trends in CTI and CAI, we filtered Data S1, retaining quadrats that had observation records across all six-decade time periods, grouping by taxonomic class. This was done to avoid the influence of the appearance or disappearance of quadrats from the dataset at different time points, which could bias estimates. After filtering Data S1, we then calculated global average CTI and CAI, including all taxonomic classes (Figure 2 C,D), as well as for each taxonomic class separately (Figure 3 A,B). We also calculated standard errors for all cases, and average CTI and CAI for each ecoregion (Figs. S2,S5). To generate estimates of $T_{max}$, AI, CTI and CAI trends for each ecoregion, we extracted the estimated marginal mean estimates after regressing the respective climate trend by ecoregion (Figure 5 A,B).

Finally, to create maps of $T_{max}$, AI, CTI and CAI trends, we averaged trends across all taxonomic classes for each quadrat (Figure 4 A-D). To generate a lag metric for CTI, we subtracted decadal trends in $T_{max}$ from decadal $CTI_{max}$ trends for each quadrat (Figure 4E). We then grouped each quadrat based on the country and averaged the lag metric. This provided average estimates of how closely turnover in communities of species tracked climate change at the country level (Figure 5D). We did not generate a lag metric for $CAI_{min}$ since the temporal trends were magnitudes smaller than AI trends and resulted in a lag metric that was almost perfectly correlated with AI trends. Finally, for each quadrat, we calculated the percentage of present species whose $STI_{max}$ value was below the maximum temperature value recorded in the most recent time period (Figure 4F). We additionally grouped all quadrats by country and calculated the average percentage of species with $STI_{max}$ values below maximum temperature conditions (Figure 5D).

*3.4 Reporting of Indices*

We compared results across all analyses to identify the most consistent and informative indices for both temperature and aridity. Based on the results, $CTI_{max}$ and $CAI_{min}$ best demonstrated community changes in response to climate trends and consistently predict species turnover in alignment with expectations of how species are anticipated to respond to changes in climate. $CTI_{mean}$ showed similar patterns of community change across taxonomic groups when compared to $CTI_{max,}$ however, it did not consistently reflect differences between species turnover groups. Additionally, $CTI_{min}$ did not consistently reflect community change or species turnover differences. All CAI metrics showed variable results with respect to community turnover in response to climate change. Specifically, changes in mammal and reptile communities were reflected by changes in $CAI_{max}$ and $CAI_{mean}$, though changes in amphibian and bird communities were not. $CAL_{min}$ was informative of community changes for amphibians and mammals, but not birds or reptiles. However, differences in $SAI_{min}$ between species turnover categories were consistent for all taxonomic groups, while $SAI_{max}$ and $SAI_{mean}$ were not consistent. Given the greater level of consistency and alignment with predictions of species turnover, we therefore report the results for $CTI_{max}$ and $CAI_{min}$ here in the SI and in the main manuscript.

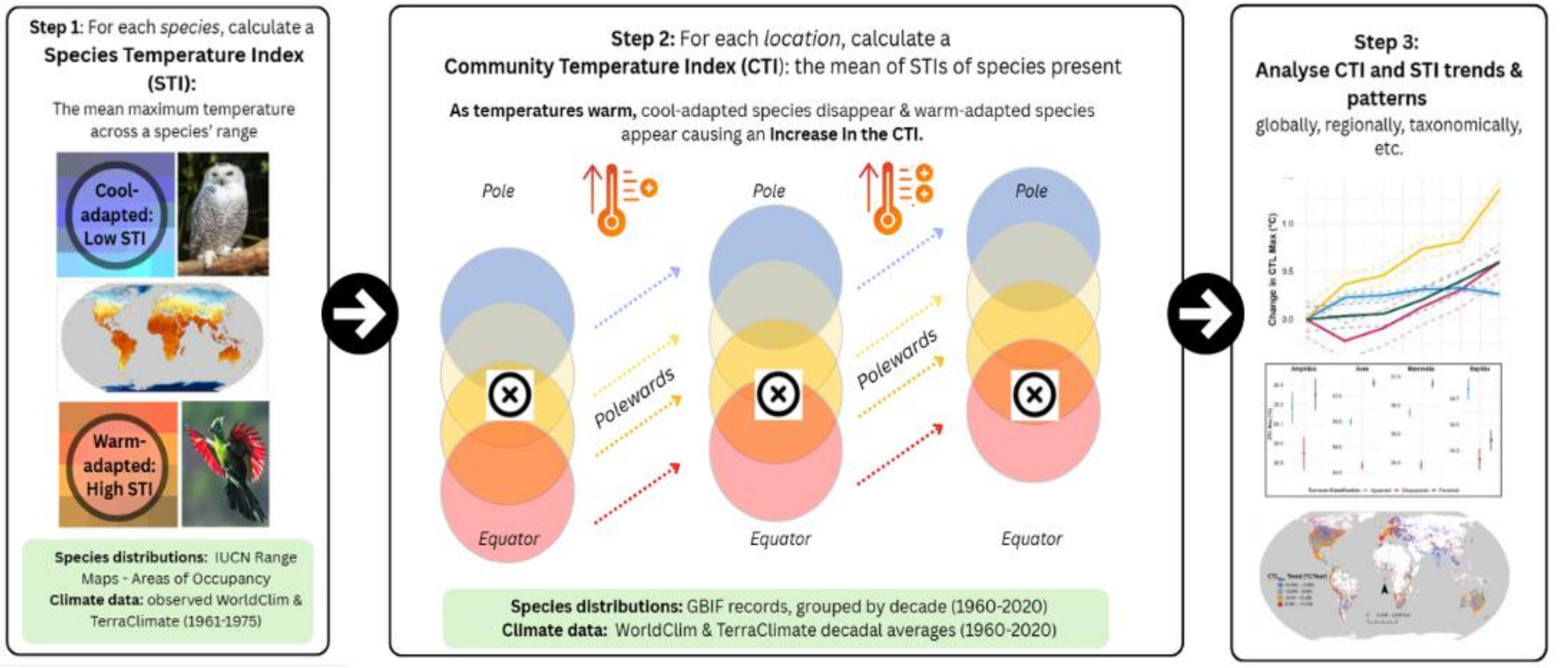

**Figure S1. Schematic diagram summarising methodological steps.** In panel 2, X indicates a location and the circles represent species with STIs varying from low (blue, cool-adapted) to high (red, warm-adapted), and panels show different time periods during which progressive warming takes place. The Species Aridity Index, Community Index and analyses were carried out in the same manner. This figure is purely illustrative and, as our study found, several areas experienced cooling and wetting.

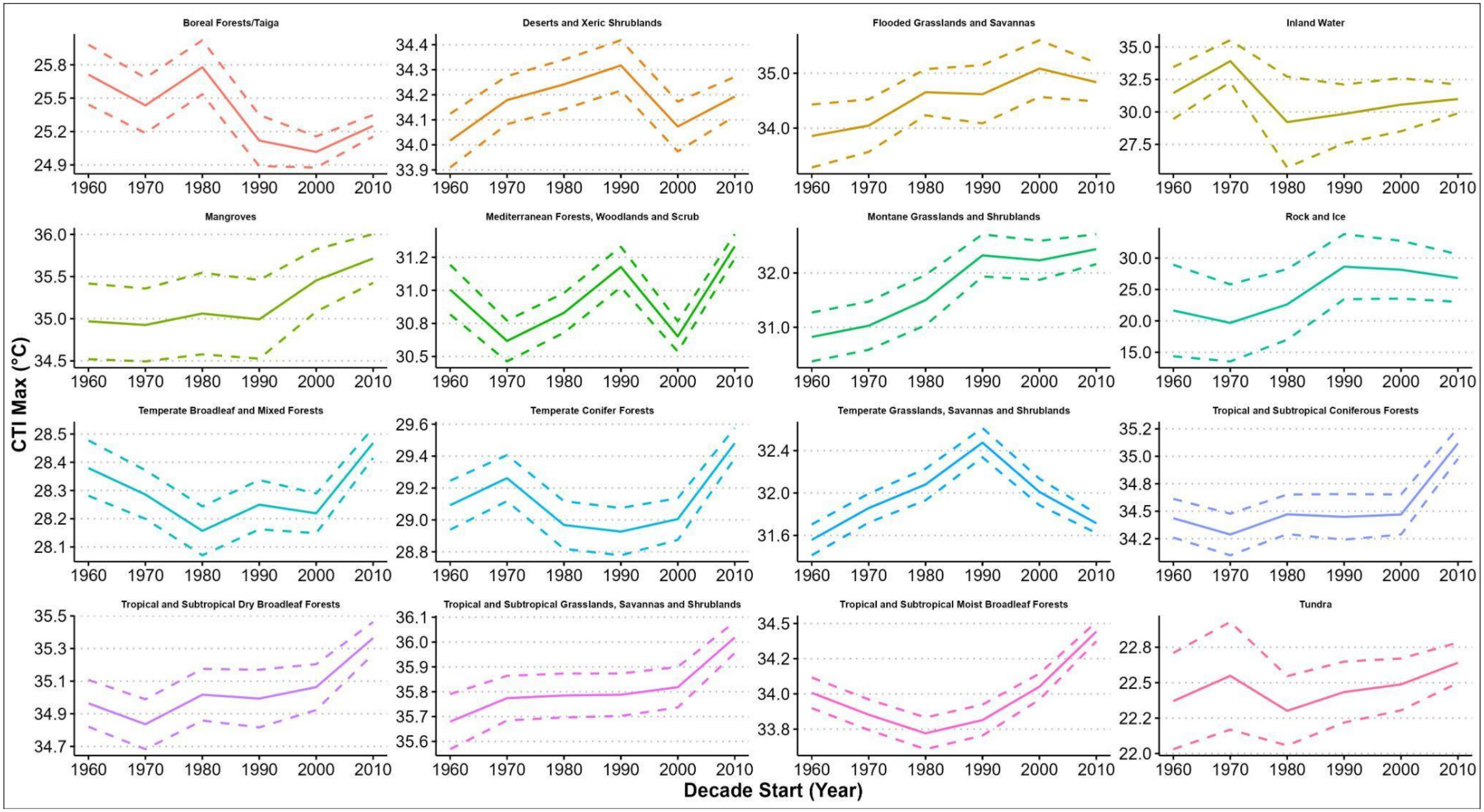

**Figure S2. Plots of global mean decadal $CTI_{max}$ by ecoregion.** Solid colored lines indicate mean trends for all aggregate taxa. Dashed lines indicate 2 standard errors.

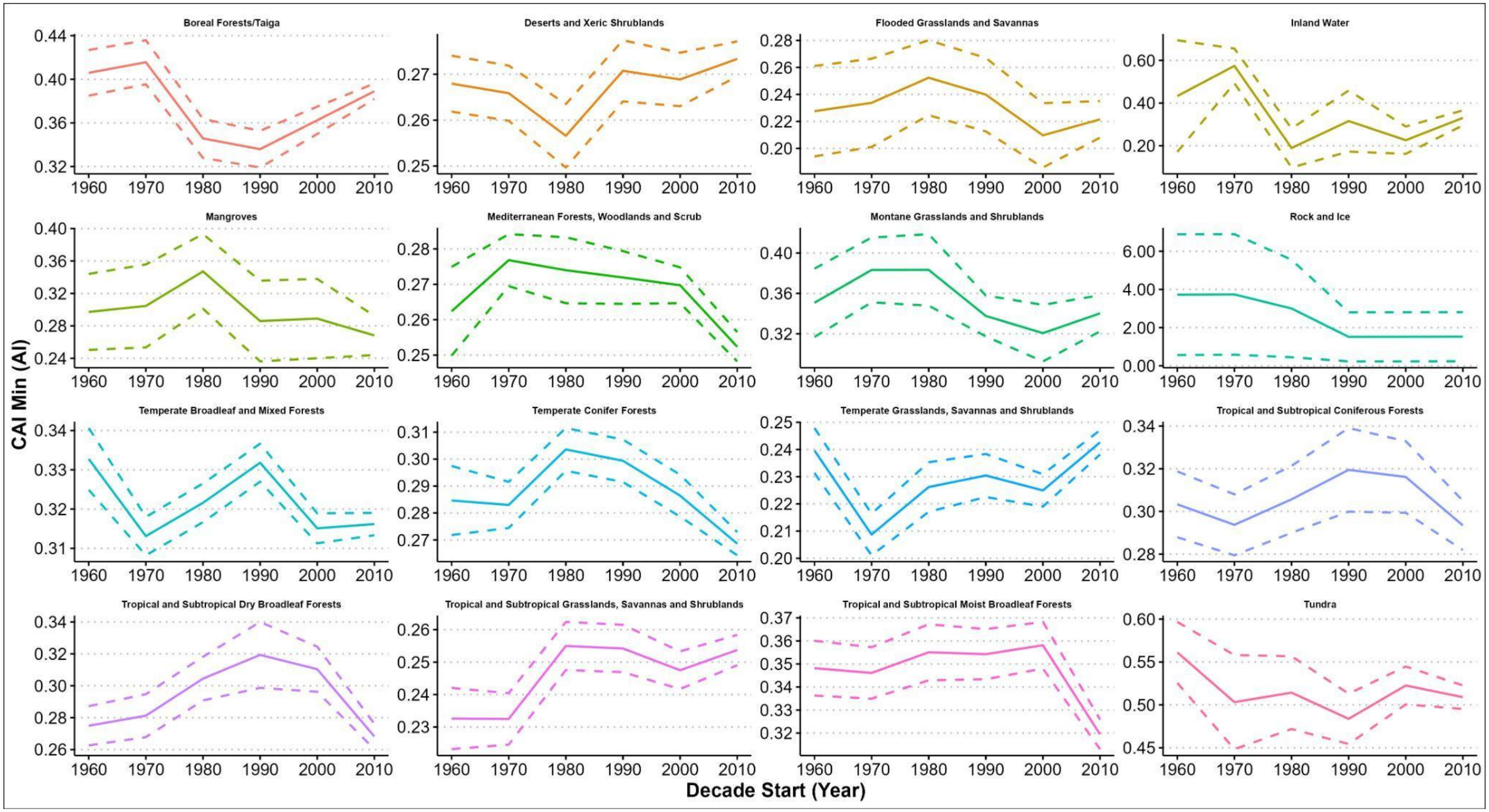

**Figure S3. Plots of global mean decadal $CAI_{min}$ by ecoregion.** Solid colored lines indicate mean trends for all aggregate taxa. Dashed lines indicate 2 standard errors.

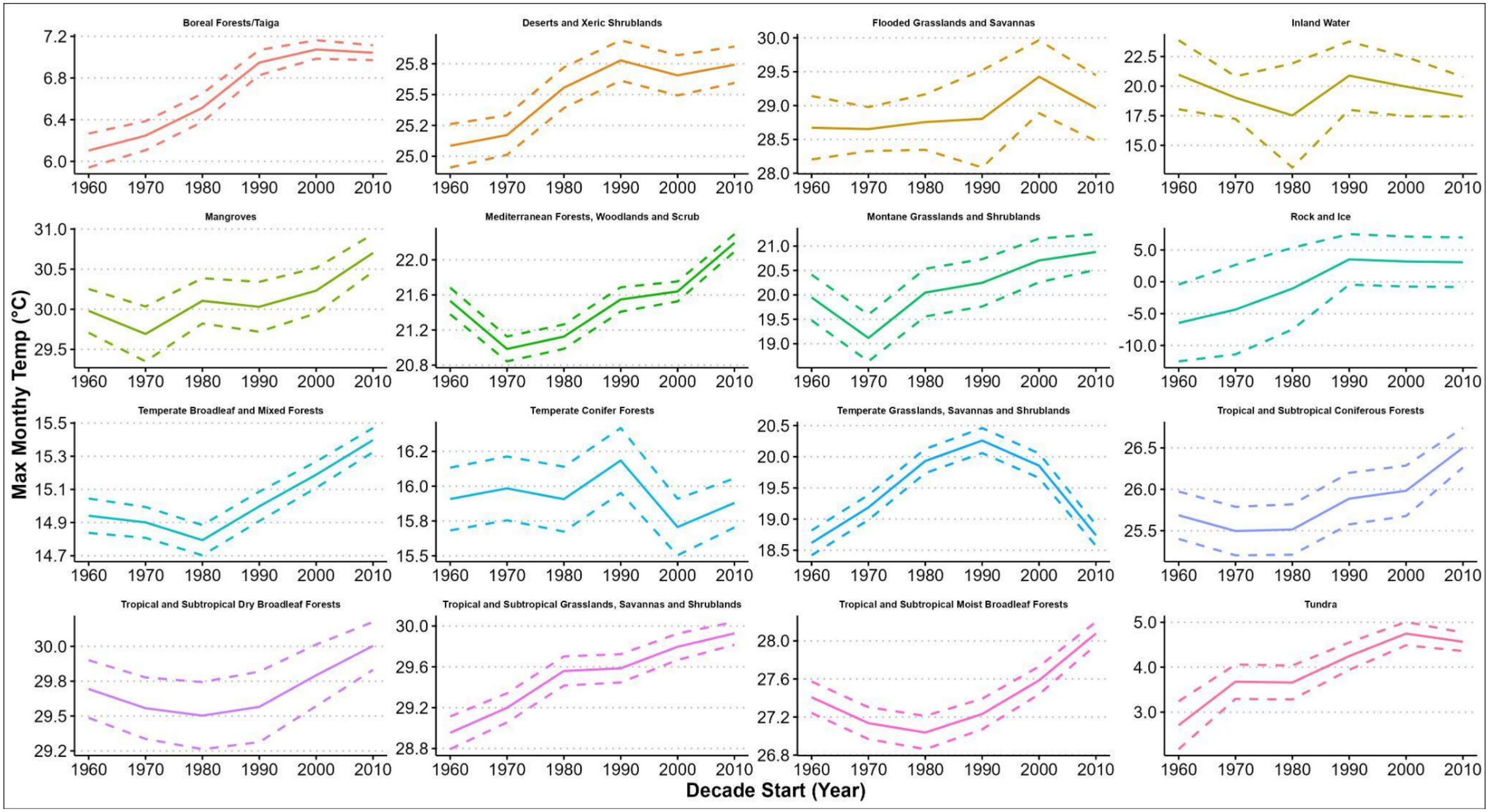

**Figure S4. Plots of global mean decadal $T_{max}$ by ecoregion.** Solid colored lines indicate mean trends for all aggregate taxa. Dashed lines indicate 2 standard errors.

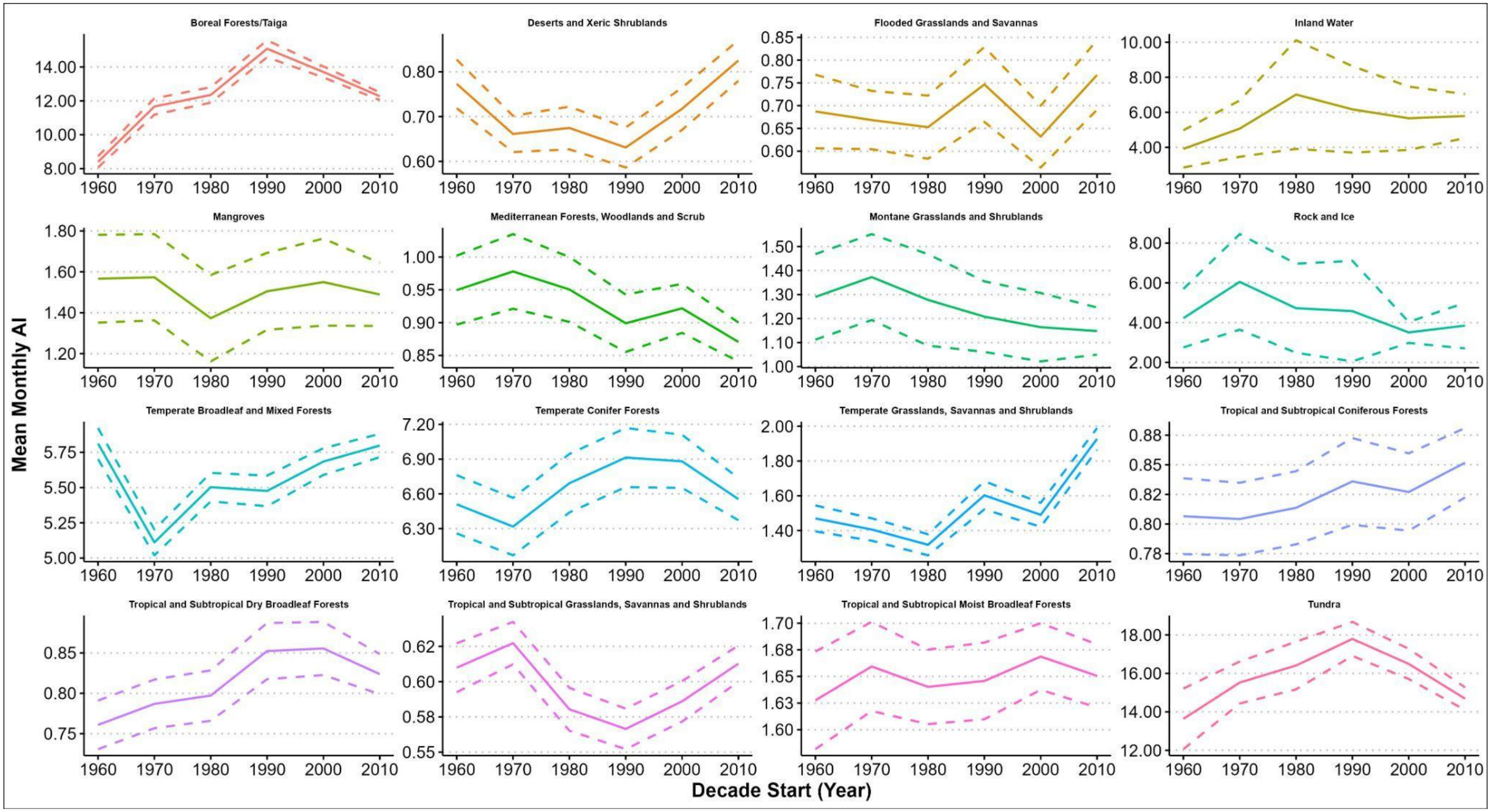

**Figure S5. Plots of global mean decadal Aridity Index by ecoregion.** Solid colored lines indicate mean trends for all aggregate taxa. Dashed lines indicate 2 standard errors.

**Table S1. Linear mixed effect model results for $CTI_{max}$ trend response to climate change.** Model coefficients for the impacts of climate change on community upper thermal limit. All continuous variables were z-score transformed before model fitting. 25,315 total observations. Marginal/Conditional $R^2 = 0.108/0.152$. Proportion of residual variation explained by fixed effects = 0.113 (marginal/1 − (conditional − marginal). All class model coefficients are presented as difference relative to reference class (Amphibia). Variable adjusted GVIF: $T_{max}$ Trend = 2.78, Class = 1.01, Elevation = 1.10, Latitude = 1.09, $T_{max}$ Trend x Class = 1.41, $T_{max}$ Trend x Elevation = 1.11, $T_{max}$ Trend x Latitude = 1.08.

max

***Fixed Effects***

| *Variable* | *Estimate* | *Std. Err.* | *p value* |
|---|---|---|---|
| **Intercept (Class: Amphibia)** | **0.1387** | **0.0450** | **0.008** |
| **Class: Aves** | **-0.3529** | **0.0174** | **< 0.001** |
| **Class: Mammalia** | **0.2216** | **0.0213** | **< 0.001** |
| Class: Reptilia | -0.0197 | 0.0212 | 0.352 |
| **$T_{max}$ trend (Class: Amphibia)** | **0.0361** | **0.0162** | **0.025** |
| Elevation | 0.0051 | 0.0064 | 0.423 |
| **Latitude (polynomial = 1)** | **-0.0527** | **0.0087** | **< 0.001** |
| Latitude (polynomial = 2) | 0.0082 | 0.0094 | 0.384 |
| $T_{max}$ trend x Class: Aves | -0.0288 | 0.0171 | 0.093 |
| **$T_{max}$ trend x Class: Mammalia** | **0.1118** | **0.0216** | **< 0.001** |
| $T_{max}$ trend x Class: Reptilia | -0.0012 | 0.0216 | 0.956 |
| **$T_{max}$ trend x Elevation** | **-0.0366** | **0.0049** | **< 0.001** |
| **$T_{max}$ trend x Latitude (polynomial = 1)** | **0.0312** | **0.0059** | **< 0.001** |
| $T_{max}$ trend x Latitude (polynomial = 2) | -0.0108 | 0.0059 | 0.069 |

***Random Effects***

| *Group* | *N groups* | *Term* | *Variance* | *Std. Dev.* |
|---|---|---|---|---|
| Ecoregion | 16 | Intercept | 0.0249 | 0.1578 |
| Residual | | | 0.4840 | 0.6957 |

**Table S2. Linear mixed effect model results for $CAI_{min}$ trend response to climate change.** Model coefficients for the impacts of climate change on the community's lower aridity limit. 30,443 total observations. All continuous variables were z-score transformed prior to model fitting. Marginal/Conditional $R^2$ = 0.007/0.020. Proportion of residual variation explained by fixed effects = 0.007 (marginal/1 − (conditional − marginal). All class model coefficients are presented as difference relative to reference class (Amphibia). Variable adjusted GVIF: AI Trend = 4.61, Class = 1.02, Elevation = 1.09, Latitude = 1.10, AI trend x Class = 1.69, AI trend x Elevation = 1.19, AI Trend x Latitude = 1.55.

| Fixed Effects | | | |
|---|---|---|---|
| ***Variable*** | ***Estimate*** | ***Std. Err.*** | ***p value*** |
| Intercept (Class: Amphibia) | -0.0759 | 0.0368 | 0.052 |
| **Class: Aves** | **0.1012** | **0.0180** | **< 0.001** |
| Class: Mammalia | -0.0201 | 0.0198 | 0.310 |
| Class: Reptilia | 0.0013 | 0.0209 | 0.952 |
| **AI trend (Class: Amphibia)** | **0.0680** | **0.0235** | **0.004** |
| **Elevation** | **0.0346** | **0.0050** | **< 0.001** |
| **Latitude (polynomial = 1)** | **0.0232** | **0.0072** | **0.001** |
| **Latitude (polynomial = 2)** | **0.0210** | **0.0072** | **0.003** |
| **AI trend x Class: Aves** | **-0.0491** | **0.0230** | **0.033** |
| AI trend x Class: Mammalia | 0.0288 | 0.0247 | 0.243 |
| **AI trend x Class: Reptilia** | **-0.0765** | **0.0299** | **0.011** |
| **AI trend x Elevation** | **-0.0281** | **0.0071** | **0.001** |
| **AI trend x Latitude (polynomial = 1)** | **-0.0248** | **0.0100** | **0.013** |
| **AI trend x Latitude (polynomial = 2)** | **-0.0218** | **0.0046** | **< 0.001** |

| Random Effects | | | | |
|---|---|---|---|---|
| ***Group*** | ***N groups*** | ***Term*** | ***Variance*** | ***Std. Dev.*** |
| Ecoregion | 16 | Intercept | 0.0149 | 0.1222 |
| Residual | | | 1.1042 | 1.0508 |

**Table S3. Linear mixed effect model results determining if $STI_{max}$ differ between turnover classifications**. Model coefficients for the impacts of turnover classification on $STI_{max}$ for all classes of terrestrial vertebrates. All class model coefficients are presented as difference relative to reference class (Amphibia). All turnover classification model coefficients are presented as difference relative to reference turnover classification (Appeared). LMMs include 707,494 total observations. $CTI_{max}$ LMM Marginal/Conditional $R^2 = 0.009/0.011$. All variables have adjusted generalised variance inflation factors ≤ 3.2.

| Variable | *Est* | *SE* | *p* |
|---|---|---|---|
| **Intercept (TC: Appeared x Amphibia)** | **30.5900** | **0.0422** | **< 0.001** |
| **TC: Appeared x Aves** | **7.3110** | **0.0160** | **< 0.001** |
| **TC: Appeared x Mammalia** | **7.1860** | **0.0167** | **< 0.001** |
| **TC: Appeared x Reptilia** | **5.5910** | **0.0170** | **< 0.001** |
| **TC: Disappeared (Amphibia)** | **-0.3708** | **0.0266** | **< 0.001** |
| **TC: Disappeared x Aves** | **-0.6268** | **0.0263** | **< 0.001** |
| **TC: Disappeared x Mammalia** | **-0.6797** | **0.0280** | **< 0.001** |
| **TC: Disappeared x Reptilia** | **-0.3064** | **0.0292** | **< 0.001** |
| **TC: Persisted (Amphibia)** | **0.2684** | **0.0236** | **< 0.001** |
| **TC: Persisted x Aves** | **0.6978** | **0.0232** | **< 0.001** |
| **TC: Persisted x Mammalia** | **0.4441** | **0.0244** | **< 0.001** |
| **TC: Persisted x Reptilia** | **-0.4625** | **0.0248** | **< 0.001** |
| **Elevation (TC: Appeared)** | **-0.0004** | **0.0001** | **< 0.001** |
| Elevation x TC: Disappeared) | 0.0001 | 0.0001 | 0.430 |
| **Elevation x TC: Persisted)** | **-0.0002** | **0.0001** | **< 0.001** |
| **Latitude (polynomial = 1) (TC: Appeared)** | **0.0094** | **0.0005** | **< 0.001** |
| **Latitude (polynomial = 2) (TC: Appeared)** | **-0.0031** | **0.0000** | **< 0.001** |
| **Latitude (polynomial = 1) x TC: Disappeared** | **-0.0023** | **0.0004** | **< 0.001** |
| **Latitude (polynomial = 2) x TC: Disappeared** | **0.0004** | **0.0000** | **< 0.001** |
| Latitude (polynomial = 1) x TC: Persisted | 0.0001 | 0.0004 | 0.905 |
| **Latitude (polynomial = 2) x TC: Persisted** | **-0.0002** | **0.0001** | **< 0.001** |

| **Random Effects** | | | |
|---|---|---|---|
| *Group* | *Term* | *Variance* | *Std. Dev.* |
| Quadrat (n = 47,542) | Intercept | 2.5960 | 1.6113 |
| Decade (n = 6) | Intercept | 0.0068 | 0.0822 |
| Residual | | 1953.0000 | 44.1895 |

**Table S4. Linear mixed effect model results determining if $SAI_{min}$ differ between turnover classifications**. Model coefficients for the impacts of turnover classification on $SAI_{min}$ for all classes of terrestrial vertebrates. All class model coefficients are presented as differences relative to the reference class (Amphibia). All turnover classification model coefficients are presented as differences relative to the reference turnover classification (Appeared). LMMs include 707,494 total observations. $CAI_{min}$ LMM Marginal/Conditional $R^2 = 0.007/0.007$. All variables have adjusted generalised variance inflation factors ≤ 3.2.

| Variable | *Est* | *SE* | *p* |
|---|---|---|---|
| **Intercept (TC: Appeared x Amphibia)** | **0.8591** | **0.0016** | **< 0.001** |
| **TC: Appeared x Aves** | **-0.7873** | **0.0013** | **< 0.001** |
| **TC: Appeared x Mammalia** | **-0.8214** | **0.0013** | **< 0.001** |
| **TC: Appeared x Reptilia** | **-0.7837** | **0.0014** | **< 0.001** |
| **TC: Disappeared (Amphibia)** | **0.0107** | **0.0021** | **0.024** |
| **TC: Disappeared x Aves** | **0.0053** | **0.0021** | **0.012** |
| **TC: Disappeared x Mammalia** | **0.0065** | **0.0023** | **0.004** |
| **TC: Disappeared x Reptilia** | **0.0099** | **0.0023** | **< 0.001** |
| TC: Persisted (Amphibia) | -0.0008 | 0.0019 | 0.663 |
| **TC: Persisted x Aves** | **-0.0299** | **0.0019** | **< 0.001** |
| **TC: Persisted x Mammalia** | **-0.0160** | **0.0020** | **< 0.001** |
| **TC: Persisted x Reptilia** | **-0.0125** | **0.0020** | **< 0.001** |
| **Elevation (TC: Appeared)** | **0.0001** | **0.0001** | **< 0.001** |
| Elevation x TC: Disappeared) | 0.0001 | 0.0001 | 0.093 |
| **Elevation x TC: Persisted)** | **0.0001** | **0.0001** | **< 0.001** |
| Latitude (polynomial = 1) (TC: Appeared) | 0.0001 | 0.0001 | 0.582 |
| **Latitude (polynomial = 2) (TC: Appeared)** | **0.0001** | **0.0001** | **< 0.001** |
| **Latitude (polynomial = 1) x TC: Disappeared** | **0.0003** | **0.0001** | **< 0.001** |
| **Latitude (polynomial = 2) x TC: Disappeared** | **0.0001** | **0.0001** | **< 0.001** |
| **Latitude (polynomial = 1) x TC: Persisted** | **-0.0003** | **0.0001** | **< 0.001** |
| **Latitude (polynomial = 2) x TC: Persisted** | **0.0001** | **0.0001** | **< 0.001** |

**Random Effects**

| *Group* | *Term* | *Variance* | *Std. Dev.* |
|---|---|---|---|
| Quadrat (n = 47,542) | Intercept | 0.0044 | 0.0662 |
| Decade (n = 6) | Intercept | 0.0001 | 0.0009 |
| Residual | | 1.2650 | 3.5571 |

**Table S5. Estimated marginal mean (EMM) of slope coefficients for community temperature and aridity indicators' response to climate change by taxonomic class.** EMM were calculated for each taxonomic class for tables S2-S3. Estimate columns indicate the number of SD change in the CTI/CAI indicator based on a 1 SD change in the indicated climate variable. Significance is indicated by 95% confidence intervals (95% CI) not overlapping 0.

| | **$CTI_{Max}$ trend ~ $T_{Max}$ trend** | | |
|---|---|---|---|
| ***Class*** | ***Estimate*** | ***SE*** | ***95% CI*** |
| Amphibia | 0.0361 | 0.0162 | 0.0045 - 0.0678 |
| Aves | 0.0073 | 0.0076 | -0.0076 - 0.0222 |
| Mammalia | 0.1479 | 0.0155 | 0.1176 - 0.1783 |
| Reptilia | 0.0349 | 0.0154 | 0.0048 - 0.0651 |
| | **$CAI_{Min}$ trend ~ AI trend** | | |
| ***Class*** | ***Estimate*** | ***SE*** | ***95% CI*** |
| Amphibia | 0.0680 | 0.0235 | 0.0219 - 0.1140 |
| Aves | 0.0189 | 0.0113 | -0.0032 - 0.0410 |
| Mammalia | 0.0968 | 0.0148 | 0.0679 - 0.1257 |
| Reptilia | -0.0086 | 0.0223 | -0.0523 - 0.0352 |

**Table S6. Estimated marginal mean (EMM) of differences in STI between turnover classifications.** EMM were calculated for each pairwise comparison between turnover categories based on the results in Table S4. Estimate columns indicate the difference in STI values between the two listed turnover groups.

| | | | | |
|---|---|---|---|---|
| | | **Amphibians** | | |
| ***Comparison*** | ***Estimate*** | ***SE*** | ***z*** | ***p*** |
| Appeared - Disappeared | 0.2346 | 0.0242 | 9.699 | < 0.001 |
| Appeared - Persisted | -0.0678 | 0.0210 | -3.221 | 0.004 |
| Disappeared - Persisted | -0.3024 | 0.0249 | -12.158 | < 0.001 |
| | | **Birds** | | |
| ***Comparison*** | ***Estimate*** | ***SE*** | ***z*** | ***p*** |
| Appeared - Disappeared | 0.8614 | 0.0193 | 44.730 | < 0.001 |
| Appeared - Persisted | -0.7656 | 0.0187 | -40.997 | < 0.001 |
| Disappeared - Persisted | -1.6270 | 0.0199 | -81.607 | < 0.001 |
| | | **Mammals** | | |
| ***Comparison*** | ***Estimate*** | ***SE*** | ***z*** | ***p*** |
| Appeared - Disappeared | 0.9142 | 0.0218 | 42.020 | < 0.001 |
| Appeared - Persisted | -0.5118 | 0.0203 | -25.273 | < 0.001 |
| Disappeared - Persisted | -1.4261 | 0.0226 | -62.979 | < 0.001 |
| | | **Reptiles** | | |
| ***Comparison*** | ***Estimate*** | ***SE*** | ***z*** | ***p*** |
| Appeared - Disappeared | 0.5410 | 0.0223 | 24.218 | < 0.001 |
| Appeared - Persisted | 0.3947 | 0.0200 | 19.699 | < 0.001 |
| Disappeared - Persisted | -0.1463 | 0.0231 | -6.347 | < 0.001 |

**Table S7. Estimated marginal mean (EMM) of differences in SAI between turnover classifications.** EMM were calculated for each pairwise comparison between turnover categories based on the results in Table S5. Estimate columns indicate the difference in SAI values between the two listed turnover groups.

| | **Amphibians** | | | |
|---|---|---|---|---|
| ***Comparison*** | ***Estimate*** | ***SE*** | ***z*** | ***p*** |
| Appeared - Disappeared | -0.01377 | 0.00194 | -7.095 | < 0.001 |
| Appeared - Persisted | -0.00064 | 0.00169 | -0.377 | 0.925 |
| Disappeared - Persisted | 0.013134 | 0.002 | 6.574 | < 0.001 |
| | **Birds** | | | |
| ***Comparison*** | ***Estimate*** | ***SE*** | ***z*** | ***p*** |
| Appeared - Disappeared | -0.0191 | 0.0016 | -12.341 | < 0.001 |
| Appeared - Persisted | 0.0292 | 0.0015 | 19.502 | < 0.001 |
| Disappeared - Persisted | 0.0483 | 0.0016 | 30.135 | < 0.001 |
| | **Mammals** | | | |
| ***Comparison*** | ***Estimate*** | ***SE*** | ***z*** | ***p*** |
| Appeared - Disappeared | -0.0203 | 0.0018 | -11.626 | < 0.001 |
| Appeared - Persisted | 0.0154 | 0.0016 | 9.458 | < 0.001 |
| Disappeared - Persisted | 0.0357 | 0.0018 | 19.609 | < 0.001 |
| | **Reptiles** | | | |
| ***Comparison*** | ***Estimate*** | ***SE*** | ***z*** | ***p*** |
| Appeared - Disappeared | -0.0236 | 0.0018 | -13.184 | < 0.001 |
| Appeared - Persisted | 0.0119 | 0.0016 | 7.3680 | < 0.001 |
| Disappeared - Persisted | 0.0355 | 0.0019 | 19.163 | < 0.001 |

**Table S8. Linear mixed effect model results for $CTI_{max}$ trend for ecoregions.** All continuous variables were z-score transformed prior to model fitting. All class model coefficients are presented as difference relative to reference class (Amphibia). Total observations were 25,315. Marginal/Conditional $R^2$ = 0.131/0.134.

| Variable | Estimate | SE | *P* < |
|---|---|---|---|
| **Intercept (Class: Amphibia)** | **0.306** | **0.041** | **0.001** |
| **Ecoregion: Deserts and Xeric Shrublands** | **-0.252** | **0.036** | **0.001** |
| **Ecoregion: Flooded Grasslands and Savannas** | **-0.154** | **0.071** | **0.029** |
| **Ecoregion: Inland Water** | **-0.550** | **0.137** | **0.001** |
| **Ecoregion: Mangroves** | **-0.269** | **0.105** | **0.010** |
| **Ecoregion: Mediterranean Forests, Woodlands and Scrub** | **-0.140** | **0.036** | **0.001** |
| Ecoregion: Montane Grasslands and Shrublands | -0.070 | 0.051 | 0.174 |
| **Ecoregion: Rock and Ice** | **-0.648** | **0.268** | **0.015** |
| **Ecoregion: Temperate Broadleaf and Mixed Forests** | **-0.135** | **0.028** | **0.001** |
| **Ecoregion: Temperate Conifer Forests** | **-0.020** | **0.033** | **0.542** |
| **Ecoregion: Temperate Grasslands, Savannas and Shrublands** | **-0.344** | **0.033** | **0.001** |
| **Ecoregion: Tropical and Subtropical Coniferous Forests** | **-0.202** | **0.054** | **0.001** |
| **Ecoregion: Tropical and Subtropical Dry Broadleaf Forests** | **-0.276** | **0.045** | **0.001** |
| **Ecoregion: Tropical and Subtropical Grasslands, Savannas and Shrublands** | **-0.233** | **0.040** | **0.001** |
| **Ecoregion: Tropical and Subtropical Moist Broadleaf Forests** | **-0.217** | **0.041** | **0.001** |
| **Ecoregion: Tundra** | **0.269** | **0.048** | **0.001** |
| **Tmax trend** | **0.053** | **0.017** | **0.002** |
| **Class: Aves** | **-0.352** | **0.017** | **0.001** |
| **Class: Mammalia** | **0.223** | **0.021** | **0.001** |
| Class: Reptilia | -0.019 | 0.021 | 0.361 |
| Elevation | 0.005 | 0.006 | 0.438 |
| **Latitude (polynomial = 1)** | **-0.055** | **0.014** | **0.001** |
| Latitude (polynomial = 2) | 0.004 | 0.012 | 0.761 |
| Tmax trend x Class: Aves | -0.029 | 0.017 | 0.086 |
| **Tmax trend x Class: Mammalia** | **0.112** | **0.022** | **0.001** |
| Tmax trend x Class: Reptilia | -0.002 | 0.022 | 0.930 |
| **Tmax trend x Elevation** | **-0.037** | **0.005** | **0.001** |
| **Tmax trend x Latitude (polynomial = 1)** | **0.023** | **0.007** | **0.001** |
| **Tmax trend x Latitude (polynomial = 2)** | **-0.016** | **0.008** | **0.035** |

| Random Effects | | | |
|---|---|---|---|
| ***Group*** | ***Term*** | ***Variance*** | ***Std. Dev.*** |
| Quadrat | Intercept | 0.00224 | 0.04732 |
| Residual | | 0.48211 | 0.69434 |

**Table S9. General linear model results for $CAI_{min}$ trend for ecoregions accounting.** All continuous variables were z-score transformed prior to model fitting. All class model coefficients are presented as difference relative to reference class (Amphibia). Total observations were 30,441. F-statistic = 19.85, $P < 0.001$, Multiple $R^2$: 0.01795, Adjusted $R^2$: 0.01704.

| Variable | Estimate | SE | $P <$ |
|---|---|---|---|
| **Intercept (Class: Amphibia)** | **-0.103** | **0.033** | **0.002** |
| Ecoregion: Deserts and Xeric Shrublands | 0.047 | 0.028 | 0.090 |
| Ecoregion: Flooded Grasslands and Savannas | 0.004 | 0.058 | 0.947 |
| Ecoregion: Inland Water | 0.127 | 0.117 | 0.278 |
| Ecoregion: Mangroves | -0.059 | 0.069 | 0.398 |
| Ecoregion: Mediterranean Forests, Woodlands and Scrub | 0.048 | 0.027 | 0.078 |
| Ecoregion: Montane Grasslands and Shrublands | -0.034 | 0.041 | 0.415 |
| Ecoregion: Rock and Ice | 0.343 | 0.218 | 0.115 |
| **Ecoregion: Temperate Broadleaf and Mixed Forests** | **0.080** | **0.022** | **0.001** |
| **Ecoregion: Temperate Conifer Forests** | **-0.075** | **0.026** | **0.004** |
| **Ecoregion: Temperate Grasslands, Savannas and Shrublands** | **0.087** | **0.026** | **0.001** |
| Ecoregion: Tropical and Subtropical Coniferous Forests | 0.010 | 0.044 | 0.822 |
| Ecoregion: Tropical and Subtropical Dry Broadleaf Forests | 0.009 | 0.036 | 0.812 |
| Ecoregion: Tropical and Subtropical Grasslands, Savannas and Shrublands | 0.048 | 0.032 | 0.127 |
| Ecoregion: Tropical and Subtropical Moist Broadleaf Forests | 0.046 | 0.032 | 0.159 |
| **Ecoregion: Tundra** | **-0.424** | **0.033** | **0.001** |
| **AI trend** | **0.096** | **0.024** | **0.001** |
| **Class: Aves** | **0.101** | **0.018** | **0.001** |
| **Class: Mammalia** | -0.021 | 0.020 | 0.295 |
| Class: Reptilia | 0.001 | 0.021 | 0.957 |
| Elevation | **0.034** | **0.005** | **0.001** |
| Latitude (polynomial = 1) | **0.042** | **0.011** | **0.001** |
| Latitude (polynomial = 2) | **0.030** | **0.009** | **0.001** |
| AI trend x class: Aves | **-0.050** | **0.023** | **0.031** |
| AI trend x class: Mammalia | 0.029 | 0.025 | 0.246 |
| AI trend x class: Reptilia | -0.077 | 0.030 | 0.010 |
| AI trend x Elevation | **-0.028** | **0.007** | **0.001** |
| AI trend x Latitude (polynomial = 1) | **-0.040** | **0.009** | **0.001** |
| AI trend x Latitude (polynomial = 2) | **-0.026** | **0.006** | **0.001** |

**Table S10. Taxonomic and spatial coverage of datasets.** Values presented in the table refer to the entire scope of coverage contained within our full datasets. The percentage of species sampled was evaluated relative to the number of terrestrial species in each class for which IUCN or Bird Life range maps are available.

| | | **Taxonomic:** | | | |
|---|---|---|---|---|---|
| **Class** | **N Observations** | **N Species** | **% Species** | **N Quadrats** | **N Observations** |
| Amphibia | 902,659 | 6,196 | 77.9 | 9,037 | 902,659 |
| Aves | 11,942,023 | 9,534 | 86.3 | 28,192 | 11,942,023 |
| Mammalia | 2,636,629 | 4,645 | 82.1 | 13,130 | 2,636,629 |
| Reptilia | 1,284,968 | 5,788 | 62.3 | 11,528 | 1,284,968 |

| | **Spatial:** | |
|---|---|---|
| ***Criteria*** | ***Value*** | ***Percentage*** |
| Global land area | 94,466,000 $km^2$ | 63.4% |
| Countries represented | 176 | 90.3% |

**Table S11. Monthly distribution of bird seasonal range maps.** The table includes the four life history stages classification for bird range maps and the corresponding months based on the hemisphere.

| Range Map | Northern Hemisphere Months | Southern Hemisphere Months |
|---|---|---|
| Breeding | May, Jun, Jul, Aug, Sep, | Nov, Dec, Jan, Feb, Mar |
| Non Breeding | Jan, Feb, Dec | Jun, Jul, Aug |
| Non Breeding Migration | Mar, Nov | Sep, May |
| Breeding Migration | Apr, Oct | Oct, Apr |

**Table S12. Summary of three types of relationships between the Community Climate Indices (CCIs) and climatic change, and possible explanations for these patterns.**

### A. MATCHED

| Component | Description |
|---|---|
| **What the Index shows** | **Changes in CCIs track climatic change.** |
| **Immigration** | Species **move** through the landscape to track shifting bioclimatic conditions. |
| **Local extinction** | Species track shifting bioclimatic conditions and may become **locally extinct** when their Species Thermal Index (STI) maximum is exceeded by maximum temperatures (Tmax), usually along the trailing edge of their distribution. |
| **Possible explanations** | Community turnover closely tracks climatic change, suggesting relatively effective dispersal, colonisation, local extinction and demographic responses to climatic shifts. |

### B. LAGGED

| Component | Description |
|---|---|
| **What the Index shows** | **Changes in CCIs are lower than climate change.** |
| **Immigration** | Species **move** through the landscape to track shifting bioclimatic conditions, but immigration of new species may be slower relative to the rate of climatic change. |
| **Local extinction** | **Extinction lag/debt** due to species persisting with STIs less than Tmax; low colonisation may contribute to delayed turnover. |

| | |
|---|---|
| **Possible explanations** | **Extinction lag/debt:** Species may persist temporarily despite unsuitable climatic conditions, or colonisation into newly suitable areas may be slow.<br>**Low sensitivity:** Organisms may be buffered *in situ* through microhabitat buffering, broad climatic tolerances, phenotypic plasticity, or behavioural flexibility.<br>**High adaptive capacity:** Species may reduce impacts through genetic adaptation or vertical redistribution (e.g., movement to higher elevations or deeper water).<br>**Data challenges:** Imperfect detection, spatial sampling bias, and low precision of historical records may underestimate turnover.<br>**Methodological challenges:** Other climatic drivers (e.g., seasonality or extreme events) may influence change, and species-poor quadrats may be excluded from analyses. |

### C. CONFOUNDED

| Component | Description |
|---|---|
| **What the Index shows** | **Changes in CCIs are unrelated to or greater than climatic change.** |
| **Immigration** | Immigration patterns may not reflect climatic tracking and may instead result from non-climatic processes. |
| **Local extinction** | Local extinction patterns may be driven primarily by non-climatic pressures rather than direct climatic effects. |
| **Possible explanations** | **Non-climatic drivers:** Species movement and local extinction may be driven by biotic interactions (e.g., disease, invasive species, predators) or anthropogenic pressures such as habitat loss and hunting.<br>**Data and methodological challenges**: Detection, sampling, and analyti[illegible] limitations may obscure or exaggerate relationships between climatic change and community turnover. |

**Table S13. List of references for downloads of GBIF observation records.** Provides the links

| GBIF Citations |
|---|
| GBIF.org (10 September 2024) GBIF Occurrence Download https://doi.org/10.15468/dl.rangg5 |
| GBIF.org (10 September 2024) GBIF Occurrence Download https://doi.org/10.15468/dl.cxz23r |
| GBIF.org (10 September 2024) GBIF Occurrence Download https://doi.org/10.15468/dl.vx7jr9 |
| GBIF.org (10 September 2024) GBIF Occurrence Download https://doi.org/10.15468/dl.w6pdvb |
| GBIF.org (11 September 2024) GBIF Occurrence Download https://doi.org/10.15468/dl.r6hzxf |
| GBIF.org (11 September 2024) GBIF Occurrence Download https://doi.org/10.15468/dl.efmq4t |
| GBIF.org (11 September 2024) GBIF Occurrence Download https://doi.org/10.15468/dl.7zz9xh |
| GBIF.org (11 September 2024) GBIF Occurrence Download https://doi.org/10.15468/dl.5zavrw |
| GBIF.org (11 September 2024) GBIF Occurrence Download https://doi.org/10.15468/dl.bf2w5w |
| GBIF.org (11 September 2024) GBIF Occurrence Download https://doi.org/10.15468/dl.td22kq |
| GBIF.org (11 September 2024) GBIF Occurrence Download https://doi.org/10.15468/dl.24mtv8 |
| GBIF.org (11 September 2024) GBIF Occurrence Download https://doi.org/10.15468/dl.24mtv8 |
| GBIF.org (03 October 2024) GBIF Occurrence Download https://doi.org/10.15468/dl.qgrjme |
| GBIF.org (03 October 2024) GBIF Occurrence Download https://doi.org/10.15468/dl.d8c6qk |
| GBIF.org (03 October 2024) GBIF Occurrence Download https://doi.org/10.15468/dl.s3nm3a |
| GBIF.org (03 October 2024) GBIF Occurrence Download https://doi.org/10.15468/dl.cwajw8 |
| GBIF.org (03 October 2024) GBIF Occurrence Download https://doi.org/10.15468/dl.qhm6nc |
| GBIF.org (03 October 2024) GBIF Occurrence Download https://doi.org/10.15468/dl.vh2kcq |
| GBIF.org (03 October 2024) GBIF Occurrence Download https://doi.org/10.15468/dl.389qa4 |
| GBIF.org (03 October 2024) GBIF Occurrence Download https://doi.org/10.15468/dl.2w9zfs |
| GBIF.org (03 October 2024) GBIF Occurrence Download https://doi.org/10.15468/dl.yygvwb |
| GBIF.org (04 October 2024) GBIF Occurrence Download https://doi.org/10.15468/dl.uumazf |
| GBIF.org (04 October 2024) GBIF Occurrence Download https://doi.org/10.15468/dl.mpa4dg |
| GBIF.org (04 October 2024) GBIF Occurrence Download https://doi.org/10.15468/dl.mysxbq |
| GBIF.org (04 October 2024) GBIF Occurrence Download https://doi.org/10.15468/dl.wcj8de |
| GBIF.org (04 October 2024) GBIF Occurrence Download https://doi.org/10.15468/dl.ehw7q2 |
| GBIF.org (04 October 2024) GBIF Occurrence Download https://doi.org/10.15468/dl.233y4w |
| GBIF.org (04 October 2024) GBIF Occurrence Download https://doi.org/10.15468/dl.kvdvtu |
| GBIF.org (04 October 2024) GBIF Occurrence Download https://doi.org/10.15468/dl.q3rvsm |
| GBIF.org (04 October 2024) GBIF Occurrence Download https://doi.org/10.15468/dl.afkfej |
| GBIF.org (07 October 2024) GBIF Occurrence Download https://doi.org/10.15468/dl.zckpex |
| GBIF.org (07 October 2024) GBIF Occurrence Download https://doi.org/10.15468/dl.g4rq2v |
| GBIF.org (07 October 2024) GBIF Occurrence Download https://doi.org/10.15468/dl.7sr79y |
| GBIF.org (09 October 2024) GBIF Occurrence Download https://doi.org/10.15468/dl.55gyyx |
| GBIF.org (09 October 2024) GBIF Occurrence Download https://doi.org/10.15468/dl.df46cw |
| GBIF.org (09 October 2024) GBIF Occurrence Download https://doi.org/10.15468/dl.7r36ua |
| GBIF.org (09 October 2024) GBIF Occurrence Download https://doi.org/10.15468/dl.26nygk |
| GBIF.org (09 October 2024) GBIF Occurrence Download https://doi.org/10.15468/dl.vxxmrs |
| GBIF.org (09 October 2024) GBIF Occurrence Download https://doi.org/10.15468/dl.kxmr43 |

(doi) for batches of all observation data used in the generation of analyses.

**Data S1. (separate file)**

Full GBIF observation data containing all climate, land use, and niche limit variables. Used for generating results for Analysis 2.

**Data S2. (separate file)**

GBIF quadrat data containing climate and CTL/CAL trend data. Used for the generation of results for analysis 1, as well as maps and figures.

**Data S3. (separate file)**

GBIF quadrat data containing Turnover Calculations. Used for the generation of figures showing how well communities are tracking climate change within countries.

**Data S4. (separate file)**

Country-level average lags in CTI max trends compared to Tmax trends, as well as the difference between average Tmax and CTImax. Used in the generation of Figure 5.
